# M2-polarized macrophages control LSC fate by enhancing stemness, homing, immune evasion and metabolic reprogramming

**DOI:** 10.1101/2022.05.06.489506

**Authors:** Isabel Weinhäuser, Diego A. Pereira-Martins, Luciana Y. Almeida, Jacobien R. Hilberink, Cesar Ortiz, Douglas R.A. Silveira, Lynn Quek, Cleide L. Araujo, Thiago M Bianco, Antonio Lucena-Araujo, Jose Mauricio Mota, Nienke Visser, Shanna M. Hogeling, Arjan Diepstra, Emanuele Ammatuna, Gerwin Huls, Eduardo M. Rego, Jan Jacob Schuringa

## Abstract

While it is increasingly becoming clear that cancers are a symbiosis of diverse cell types and tumor clones, the tumor microenvironment (TME) in acute myeloid leukemias (AML) remains poorly understood. Here, we uncover the functional and prognostic relevance of an M2-polarized macrophage compartment. Intra bone marrow co-injection of M2d-macrophages together with leukemic blasts that fail to engraft on their own now induce fatal leukemia in mice. Even a short-term two-day *in vitro* exposure to M2d macrophages can “train” leukemic blasts after which cells are protected against phagocytosis, display increased mitochondrial metabolism and improved *in vivo* homing, resulting in full-blown leukemia. Single-cell RNAseq analysis of AML associated macrophages revealed metabolic-related pathways such as Fatty Acid Oxidation and NAD^+^ generation as therapeutical targetable vulnerabilities. Our study provides insight into the mechanisms by which the immune landscape contributes to aggressive leukemia development and provides alternatives for effective targeting strategies.

## Introduction

AML is a complex and aggressive disease characterized by a high genetic heterogeneity and clonal diversity across patients (1–4). The expansion of AML leukemic stem cells (LSC) and their progeny rapidly usurps the bone marrow (BM) niche to impede healthy hematopoiesis, reshape BM-derived stromal cell function and escape immune surveillance (5–7). Much like healthy hematopoietic stem cells (HSC), LSCs reside in BM niches to ensure their survival, facilitate AML progression, and escape cytotoxic therapy (8–10).

Within the BM niche, macrophages can regulate the fate of HSC during homeostasis. *In vivo* macrophage depletion promotes increased mobilization of HSC (11). Moreover, Hur et al., revealed that CD234^+^ macrophages can interact with CD82^+^ long-term HSC to support quiescence in the endosteal region (12), while others identified erythroblastic island macrophages to promote erythropoiesis (13, 14). Thus, the BM harbors diverse macrophage populations, which can be exploited by LSCs. In this context, Mussai et al. detected increased Arginase 2 activity released from AML blast cells to promote M2-macrophage polarization and inhibit T-cell proliferation (15). Moreover, Al-Matary et al. observed increased infiltration of monocytes/macrophages with pro-leukemogenic functions using AML murine models and identified the transcriptional repressor *Gfi1* to regulate M2 polarization (16). Finally, a recent single cell RNA (scRNA) sequencing study revealed that the heterogeneity of AML is not only inherent to the leukemic blast cell itself but extends to the stromal and immune cells within the BM (17). By analyzing 16 AML patients the authors identified eight macrophage subpopulations with 5 displaying immunosuppressive features.

Here, we show that direct interaction with M2-polarized macrophages drives aggressive *in vivo* leukemia development of primary favorable AML and of acute promyelocytic leukemia (APL) cells, which are otherwise notoriously difficult to engraft in xenograft models. Direct interaction between M2-polarized macrophages and leukemic blasts enhances immune evasion, homing, stemness, and alters metabolism via direct exchange of mitochondria from macrophages to leukemic blasts, transforming them into more aggressive leukemic stem cells. We uncover clear heterogeneity within the macrophage landscape across AML patients which has clinical importance since the presence of M2-polarized macrophages in the BM correlates with the poorest clinical outcomes. Using single-cell RNA seq technology, we identified metabolism-related pathways that were up-regulated in AML-associated macrophages (AAM), when compared to healthy BM samples. Among these pathways, we show that targeting fatty acid oxidation (FAO) and NAD^+^ generation in AAM, decreases their capacity to support blast cells. Finally, we developed a novel biomarker panel which allows the identification of this patient group with high AAM infiltration and associates with dismal outcome.

## Results

### Heterogeneity in the macrophage landscape in AML: the presence of CD163^+^/CD206^+^ leukemia-associated macrophages identifies patients with the poorest prognosis

Using the TCGA cohort (18) in which RNA sequencing was performed on the mononuclear cell (MNC) fraction of AML patients, we observed that high expression of *CD163* predicted poor overall survival (OS) (Figure 1A). This was initially rather surprising to us, considering that CD163 is typically expressed on more mature myeloid cells, and since more committed AMLs usually associate with better survival as compared to more immature AMLs (19, 20). Therefore, we prospectively investigated whether the expression of CD163 was actually arising from the leukemic blasts themselves, or rather from myeloid committed cells that are typically also present within the MNC fraction from which RNA is oftentimes extracted for transcriptome studies. Multiparametric flow cytometry (FACS) analysis revealed that expression of CD163, along with another M2 marker CD206 and the M1 marker CD80, primarily emerge from a more mature SSC^high^/CD45^high^/HLA-DR^+^/CD14^+/-^/CD16^+/-^ myeloid subpopulation (hereafter called AAM, AML-associated macrophages) (Figure 1B-C). We previously performed transcriptome analysis on AML patient samples that were sorted into leukemic stem cell-enriched CD34^+^ populations and their non-self-renewing more committed CD34^-^ progeny (21). These sets were now used for CIBERSORT analyses (22), which showed that it was indeed the CD34^-^ fraction that was most enriched for signatures associated with more mature myeloid cells (Figure 1D). These data were further confirmed by Cellular Indexing of Transcriptomes and Epitopes by Sequencing (CITE-seq) data (Figure 1E)(23). We questioned whether the quantification of macrophages in the BM by FACS would result in an underrepresentation due to difficulties in efficiently retrieving macrophages using BM aspirates, but comparison of FACS data with immunohistochemistry staining for CD163 indicated no significant differences (Figure 1F).

**Figure 1.**
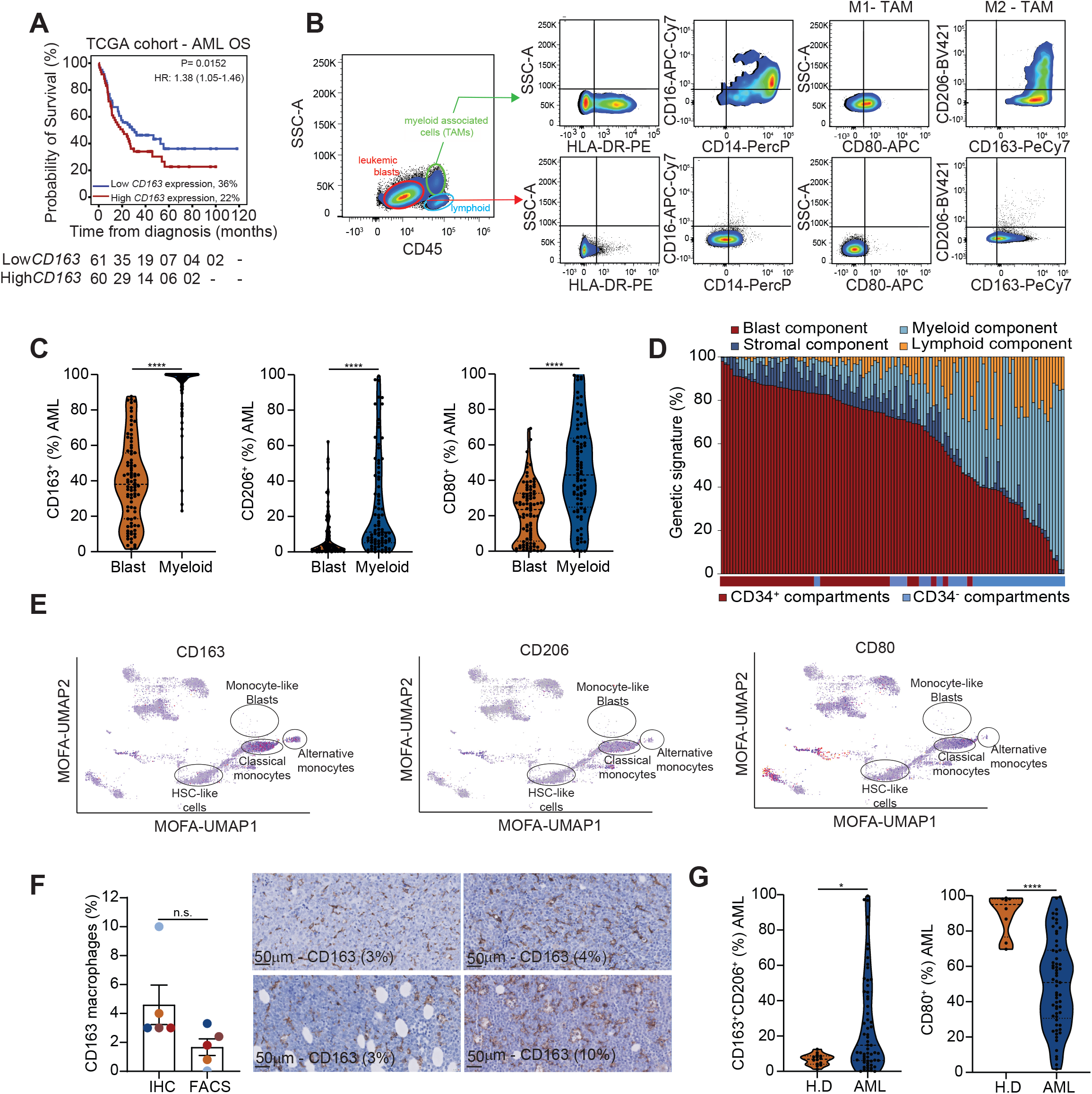
The poor prognostic value of M2 macrophages in AML. (A) Overall survival (OS) in relation to *CD163* expression in the TCGA cohort. (B) Representative FACS plot of macrophage marker expression in different AML CD45^+^ subpopulations. (C) The expression of macrophage markers in the AML blasts versus the mature myeloid population. (D) CIBERSORT analysis on the paired CD34^+^/CD34^-^ transcriptomes of AML patients. (E) UMAP data of single-cell surface protein expression of human AML patients (n = 31586 single cells) display the expression of the studied macrophage markers. (F) The amount of CD163^+^ macrophages quantified by IHC and FACS. Representative IHC pictures for CD163 in AML samples. (G) The level of M2 (CD163^+^/CD206^+^) and M1 (CD80^+^) markers measured by FACS in healthy donors (H.D., BM=5, PB=8) and AML patients (BM=33, PB=28). (A) Survival curves were compared using the log-rank test. (C and G) Wilcoxon signed rank test (2-sided) \**P*<0.05, \*\*\**P*<0.0001. (F) Mann Whitney test for unpaired data (2-sided).\**P*<0.05.

Considerable heterogeneity was observed in AAMs between individual patients, with some patients harboring predominantly tumor-suppressive M1 macrophages, while others predominantly displayed tumor-supportive M2 macrophages or both (Figure 1G, supplemental Figure 1A). Furthermore, we observed that patients with a high M2 macrophage content were more often carrying the *FLT3*-ITD mutation and more likely to be categorized within the intermediate/adverse risk group (defined by the European Leukemia Net - ELN) than patients with low M2 macrophage content or high M1 content (supplemental Figure 1B). Our data shows at the protein level, that the poor OS prediction associated with high CD163/CD206 mRNA expression (24, 25) is not necessarily driven by intrinsic AML biology but could be attributed to the presence of an AML tumor supportive niche.

### M2d macrophages support *in vivo* engraftment and leukemogenesis

Next, we evaluated whether M2 macrophages could support leukemogenesis *in vivo*. As proof of concept, we started with primary Acute Promyelocytic Leukemia (APL) samples, a subgroup of AML characterized by the *PML-RARA* translocation occurring in 95% of the patients (26) and known to be particularly difficult to engraft (27). In a first set of experiments, we injected human peripheral blood (PB) derived nonpolarized (M0) and M2d (IL-6 polarized) macrophages into the BM of NSGS mice followed by the transplant of primary APL cells via the retro-orbital route. Control mice only received primary APL cells (Figure 2A). As expected, none of the mice receiving APL blasts without macrophages succumbed to leukemia (Figure 2B-E). In stark contrast, at week twelve post-transplant mice injected with M0 and M2d macrophages presented leukocytosis indicating the manifestation of leukemia (Figure 2B). Mice were sacrificed to determine the level of human APL blasts in each leg and spine (as an internal control). The percentage of human CD45^+^ cells was increased in macrophage-injected mice compared to controls (Figure 2C). Total human APL blast cells were substantially increased for M0/M2d legs when compared to control and was superior in the presence of M2d compared to M0 macrophages (Figure 2C). Engrafted BM cells displayed human promyelocyte characteristics (Figure 2D). In line with leukemia onset, macrophage recipient mice displayed increased spleen weight (Figure 2E).

**Figure 2.**
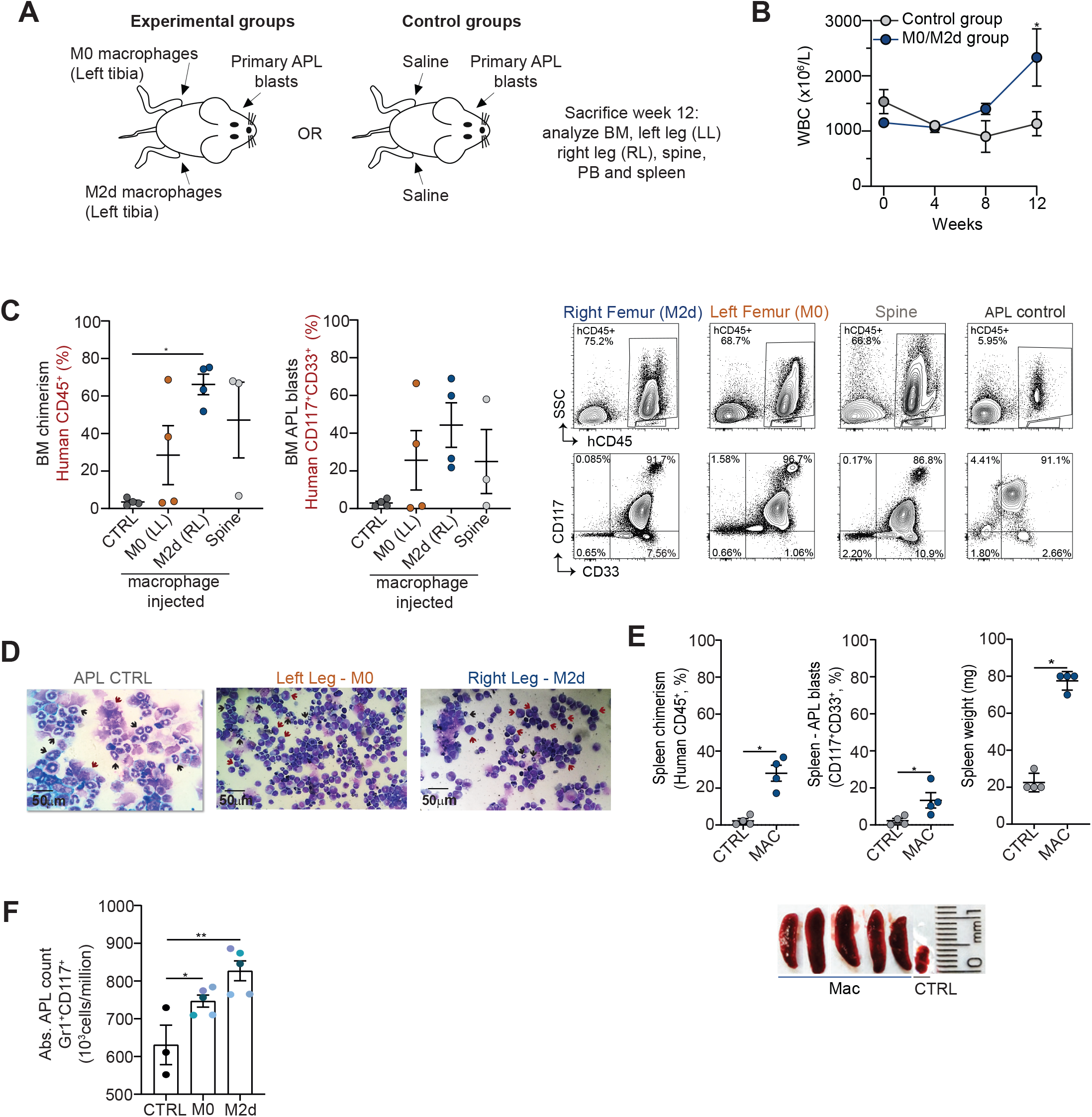
M2d macrophages support *in vivo* engraftment and leukemogenesis. (A) Experimental setup. (B) Leukocyte count in transplanted mice. Each dot represents the mean of 6 independent APL patients. (C) Bar graphs and representative FACS plots of the percentage of human CD45^+^ only (left panel) and human APL blast cells (CD117^+^/CD33^+^) detected in the BM of mice without (control) or with co-injected macrophages (right panel) (N=4). LL=Left Leg, injected with M0 macrophages; RL=Right Leg, injected with M2d macrophages. The spine was also analyzed from mice in which the macrophages were injected into the tibia. Representative FACS plot (D) Representative cytospins of the BM of mice transplanted with APL blasts and co-injected M0 or M2d macrophages. Arrows indicate human APL blast cells. (E) Human CD45^+^ and APL blasts (CD117^+^/CD33^+^) cells (%) detected in the spleen (N=4) and spleen weight. Representative spleen pictures of mice injected without (control) and with macrophages (Mac). (F) Bar graph of absolute murine Gr1^+^CD117^+^ APL blast counts detected in the BM of mice without (control) or with co-injected macrophages (Mac) 10 days post-transplant. Two-way (B) analysis of variance (ANOVA). (C and F) Kruskal-Wallis test. \**P*<0.05. (E) Mann Whitney test for unpaired data (2-sided). \**P*<0.05

These data were further validated in our murine *PML-RARA* model (28), whereby the injection of murine BM-derived M0 and M2d macrophage significantly increased the absolute number of murine APL blast cells retrieved from the BM compared to control mice 10 days after the transplant (Figure 2F). These data indicate that alterations in the BM niche can impact on the progression of leukemia and suggest that the presence of an immunosuppressive environment can facilitate the engraftment of favorable AML samples.

### Co-culture of primary APL patient cells on M2d macrophages allows the development of full-blown leukemia in a xenograft model

Based on the premise that macrophage-injected NSGS mice enabled APL engraftment and infiltration to distant tissues, we hypothesized that a pre-culture of APL cells on macrophages might be sufficient to alter the cell fate of LSCs and improve their engraftability and initiation of leukemia. We co-cultured primary human and murine APL cells on human PB or murine BM derived M0/M2d macrophages for 48hrs. Subsequently, a substantial fraction of leukemic blasts was phagocytosed leaving 8.3 and 10.5% of the original input of human and murine blasts, respectively (supplemental Figure 2A-B). Next, we injected 1.5×10^5^ “trained” cells retro orbitally into NSGS mice, while control mice received 1×10^6^ (corresponding to a 6.6-fold increase) non-cultured cells, or cells precultured on mesenchymal stromal cells (MSCs)(Figure 3A). Indeed, only human APL cells pre-cultured on macrophages were able to induce full-blown leukemia and cells pre-cultured on M2d macrophages consistently were more aggressive (Figure 3B-F). APL cells pre-cultured on MSCs did not induce leukemia (supplemental Figure 2C-D).

**Figure 3.**
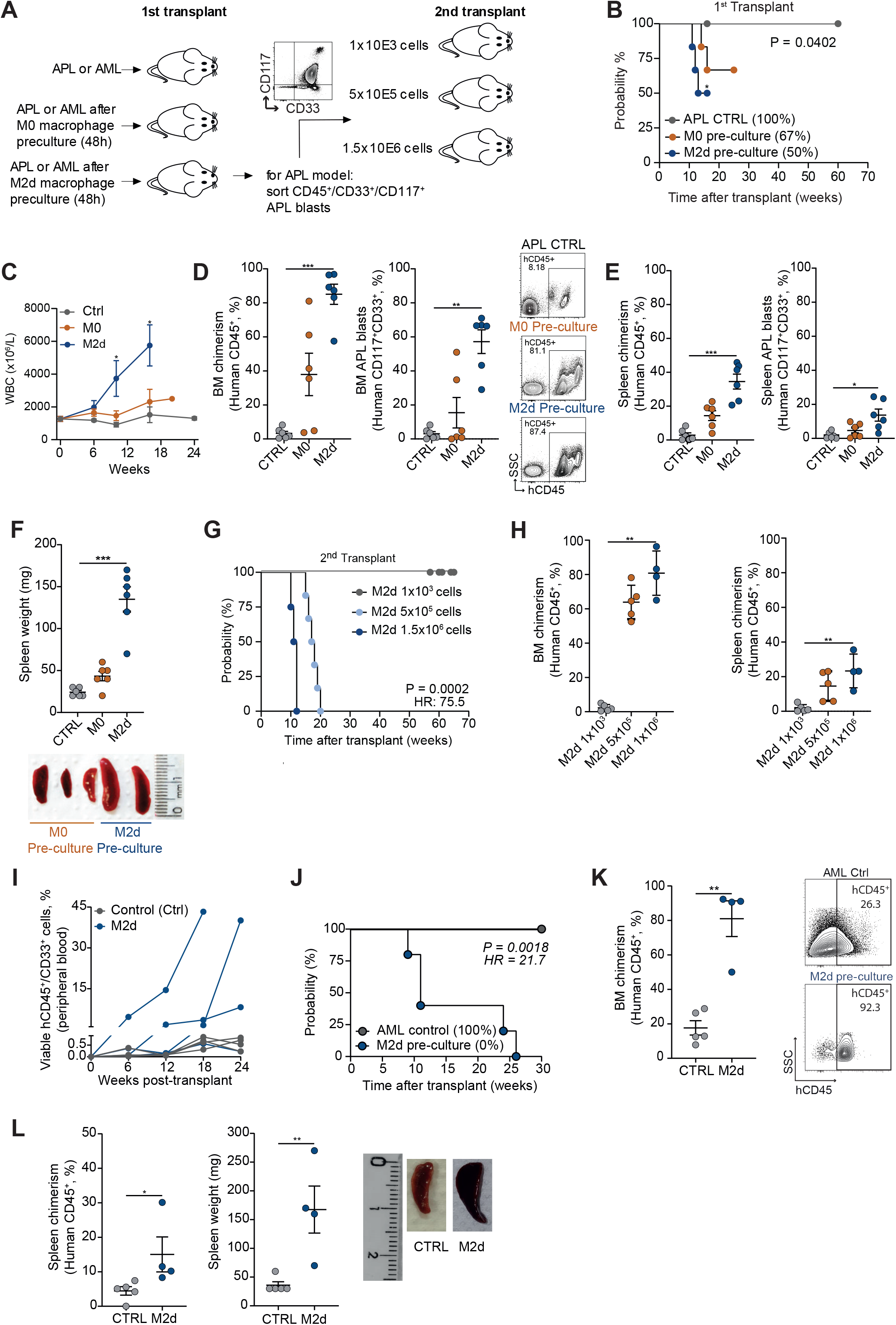
Primary APL blast cells pre-cultured on human M2d macrophages generate fatal leukemia. (A) Experimental setup. (B) OS of mice transplanted with primary APL blast transplanted without pre-culture or after pre-culture on M0 or M2d macrophages for 48h. (C-F) Mice transplanted with APL blasts without pre-culture (Ctrl) or after preculture on M0 or M2d macrophages were analyzed for WBC counts (C), for human CD45^+^/CD117^+^/CD33^+^ chimerism (%) measured in the BM (D), for human CD45^+^/CD117^+^/CD33^+^ chimerism (%) measured in the spleen (E), and spleen weight with representative spleen pictures of mice injected with M0 and M2d macrophages (F). (G) OS of secondary transplanted mice receiving 1.5×10^6^, 5×10^5^ or 1×10^3^ sorted human CD33^+^CD117^+^ APL blast cells from the primary transplant described in B-F. (N=3-6 per group). (H) Human CD45^+^ chimerism levels (%) measured in the BM and spleen of secondary transplanted mice. (I) Mice transplanted with AML blasts without pre-culture (Ctrl) or after pre-culture on M2d macrophages were analyzed for human CD45^+^/ CD33^+^ chimerism measured in the peripheral blood 6, 12, 18, and 24 weeks post-transplant. (J) OS of mice transplanted with primary AML blast transplanted without pre-culture (control) or after pre-culture on M2d macrophages for 48h. (K-L) Mice transplanted with AML blasts without pre-culture (Ctrl) or after pre-culture on M2d macrophages were analyzed for human CD45^+^ chimerism (%) measured in the BM (K), for human CD45^+^ chimerism (%) measured in the spleen and spleen weight (L). (B; G and J) Each dot represents an individual mouse OS curves were estimated using the KM method, and the log-rank test was used for comparison. Two-way (C) analysis of variance (ANOVA). *\*P* <0.05, \*\**P*<0.01, \*\*\**P*<0.001. (D-F and H) Kruskal-Wallis test. \**P* <0.05, \*\**P*<0.01, \*\*\**P*<0.001. (K-L) Wilcoxon signed rank test (2-sided). \**P*<0.05.

Next, hCD45^+^/hCD117^+^/hCD33^+^ engrafted APL blast from the primary transplanted mice were sorted and used for secondary transplant (now without “training” on macrophages) in limiting dilution (1×10^3^, 5×10^5^ and 1.5×10^6^ APL cells). After passage through primary mice the cells retained self-renewal and secondary transplanted mice developed full-blown APL when transplanted with 1.5×10^6^ (median OS 10 weeks) or 5×10^5^ cells (median OS 14 weeks) (Figure 3G-H and supplemental Figure 2E). Mice transplanted with 1×10^3^ cells only presented transient engraftment (Figure 3G-H and supplemental Figure 2F).

Similar results were obtained in our murine model, in which transgenic *PML-RARA*-expressing blast cells were transplanted into sub-lethally irradiated recipients after training on macrophages. Chimerism levels measured at week 8 indicated significantly increased APL CD45.2^+^ chimerism for mice transplanted with M0/M2d macrophage exposed murine APL cells. The median OS was 9.5 and 6.2 weeks for mice that received M0 and M2d pre-cultured cells, respectively, compared to 14 weeks for control mice (supplemental Figure 2G-H). M2d-precultured APL cells presented a more undifferentiated blast-like cell morphology compared to non-cultured APL blasts, which displayed a higher frequency of cells with cruller shaped nuclei post-sacrifice, characteristic for intermediate myeloid murine cells, which was confirmed by FACS (supplemental Figure 2I-J).

Given these results, we decided to transplant other favorable AML subtypes (n=5) such as *NPM1* and inv(16)(p13.1q22) mutated AMLs after training on M2-polarized macrophages. For all cases, pre-culturing on M2d macrophages allowed fatal leukemia development with high infiltration of leukemic blasts in the BM and spleen (Figure 3I-L).

### “Trained” AML/APL blasts are protected against phagocytosis, display improved homing capacity, and adapt a more OXPHOS-like state

Considering that a substantial fraction of leukemic cells was phagocytosed after a two-day co-culture on M2d macrophages, we questioned whether the remaining cells acquired the capacity of immune evasion to protect themselves against phagocytosis. Although NSG mice lack an adaptive immune system, they have functional macrophages with increased phagocytic activity (29), which could impede successful engraftment of primary AML cells. Moreover, MISTRG mouse models expressing the human protein SIRPα display improved engraftment in part due to avoiding phagocytosis of human cells (30). To test this hypothesis, leukemic AML blasts were co-cultured on M2d macrophages for two days after which the remaining cells were harvested and used for a new phagocytosis assay on fresh macrophages. Indeed, compared to the level of phagocytosis of uncultured cells (on average 40%) we observed a significant reduction in phagocytosis when leukemic blasts were “trained” on M2d macrophages (Figure 4A). “Don’t eat me signals” CD47 (31) or CD24 (32) were negatively correlated with AML phagocytosis (Supplemental Figure 3A-B) although no increase of these markers was observed after two days coculture on M2d macrophages (Supplemental Figure 3C). Of note, a recent paper published by Moore et al., indicated that LC3–associated phagocytosis of AML apoptotic cells by macrophages triggers the STimulator of INterferon Genes (STING) pathway thereby inhibiting AML growth (33). When we evaluated the STING genes in macrophages post-polarization and after AML co-culture, we did not observe a significant difference in expression in our setting except for *MMP9*, which was decreased (Supplemental Figure 3D). Contrarily to the “don’t eat me” signals, we noticed that surface calreticulin (CALR), an “eat me signal” predicting better prognosis in AML/APL (34) positively correlated with phagocytosis and was down-regulated upon M2d co-culture (Supplemental Figure 3B, Figure 4B). A study published by Lin et al., described Stanniocalcin 1 (STC1) to trap CALR in the mitochondria as a mechanism to reduce surface CALR expression and evade phagocytosis (35). Indeed, we find increased mRNA expression of *STC1* upon coculture with M2d macrophages (Figure 4C).

**Figure 4.**
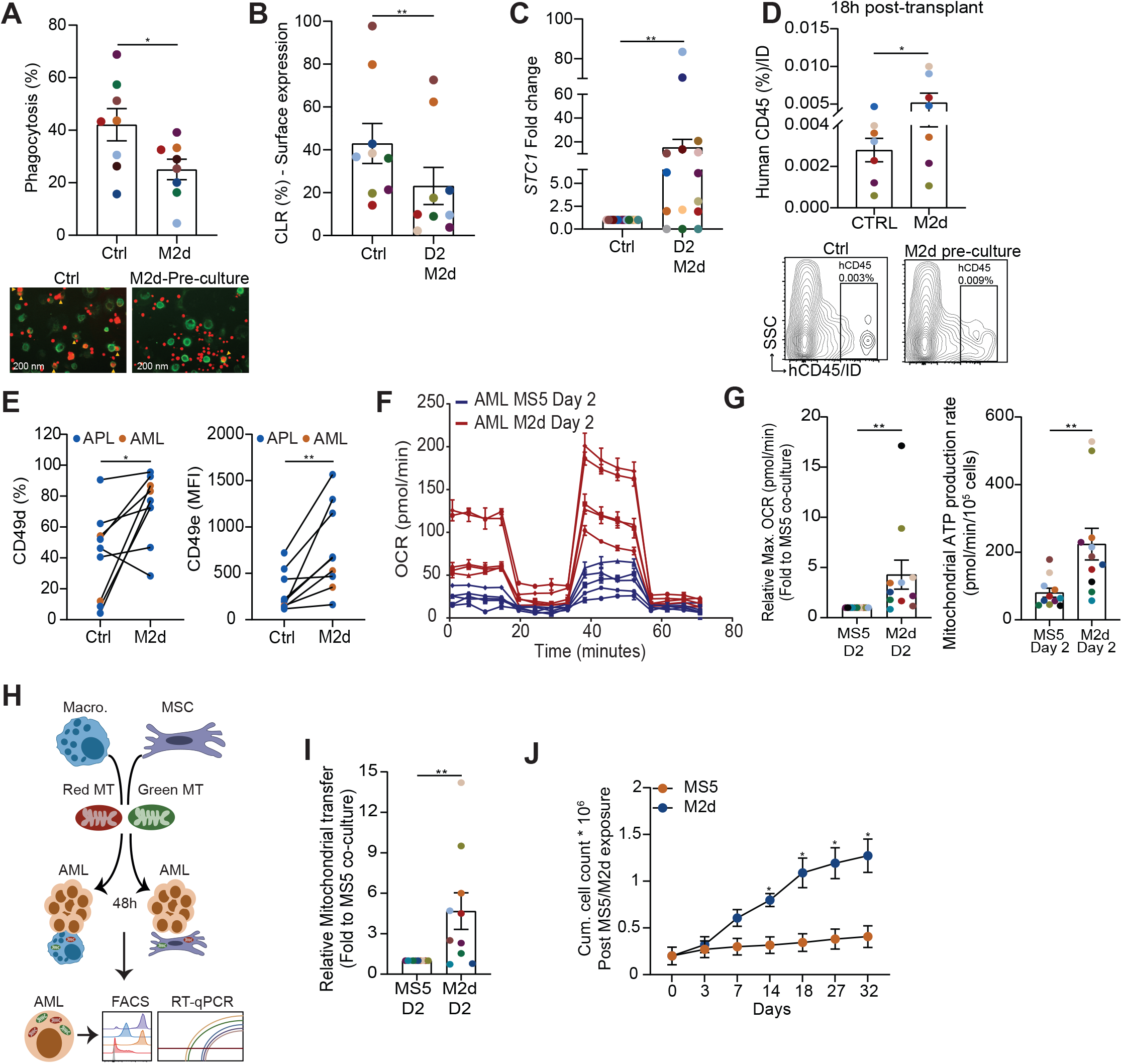
M2d macrophages reprogram primary AML cells via different biological pathways. (A) Phagocytosis of primary AML at diagnosis and after a 48h co-culture on M2d macrophages. (B) Protein expression of Calreticulin (CLR) measured on primary AML cells at diagnosis and after a 48h co-culture on M2d macrophages. (C) *STC1* gene expression measured in primary AML samples at diagnosis and after a 48h co-culture on M2d macrophages. Data is plotted as a fold change compared to diagnosis. (D) *In vivo* homing assay. Mice were injected with stained APL cells (Incucyte Dye®, ID) without prior co-culture (Control) samples or after a 48h co-culture on M2d macrophages. 18h post-transplant the levels of human CD45 (%) was analyzed in the BM. Representative FACS plots are shown. (E) CD49d (%) and CD49e (MFI) in primary APL/AML blasts at diagnosis and after a 48h co-culture on M2d macrophages. (F) Oxygen consumption rate (OCR) of primary AML blasts exposed to either MS5 or M2d macrophages for 48h. (G) Relative maximal OCR of primary AML blasts exposed to either MS5 cells or M2d macrophages. Data is plotted as a fold change compared to MS5 cells. Mitochondrial ATP production rate of primary AML blast cells exposed to either MS5 cells or M2d macrophages. (H) Experimental scheme. (I) Mitochondrial (MT) transfer measured in primary AML cells after being exposed to mitochondrial labelled MS5 and M2d cells for 48h. (J) Cumulative cell count in liquid culture of primary AML cells exposed to either MS5 or M2d macrophages for 48h. (A-G and I) each dot represents an individual patient (J) represent at least 3 biological replicates for all experiments. (A-E; G and I) Wilcoxon signed rank test (2-sided). \**P*<0.05, \*\**P*<0.01, \*\*\**P*<0.001. Two-way (J) analysis of variance (ANOVA). \**P* <0.05, \*\**P*<0.01.

One of the several hurdles AML blast cells must overcome after transplantation is to find their niche. We observed better homing of APL cells to the BM after prior co-culture on M2d macrophages compared to uncultured cells of the same patient (Figure 4D). Similar results were observed in an *in vitro* transwell migration assay (supplemental Figure 3E). Concurrently, we detected enhanced surface expression of the adhesion receptors CD49d (%) and an increase of the CD49d-f mean fluorescence intensity (MFI) upon M2d co-culture versus AML cells at diagnosis (Figure 4E and Supplemental Figure 3F). Moreover, APL blast cells harvested from primary and secondary transplants, as well as AML blasts harvested from primary transplants, displayed increased CD49d expression (supplemental Figure 3G-H).

We wondered whether coculture on M2d macrophages would impact on cellular metabolism of leukemic cells. Seahorse measurements confirmed that the functional respiration and extracellular acidification rate were enhanced in HL60 cells and primary AML samples (n=11) when co-cultured on M2d macrophages compared to MS5 controls (Figure 4F-G and supplemental Figure 4A-B). The increased basal and maximum oxygen consumption rate (OCR) suggested enhanced mitochondrial metabolism, which prompted us to determine whether macrophages, similar to MSC cells, can transfer mitochondria to primary AML cells (36) (Figure 4H). AML cell lines and primary AML cells were co-cultured for 48hrs on mitochondria-labeled M2d macrophages or MS5 cells. FACS analysis revealed an efficient mitochondrial transfer from M2d macrophages to primary AML leukemic blasts and AML cell lines, which was more efficient compared to mitochondrial exchange from MS5 cells in primary AML blasts (Figure 4I, supplemental Figure 4C). Real-time qPCR evaluation of the mitochondrial DNA content further confirmed mitochondrial transfer, which was even twice as efficient when compared to transfer from MS5 cells (supplemental Figure 4D). Finally, we assessed the long-term effects of macrophages on leukemic cells by evaluating the colony unit formation and proliferation capacity post macrophage exposure compared to MS5. Two day “training” on M2d macrophages significantly increased colony formation capacity (supplemental Figure 4E) and endowed AML cells with long term proliferation in liquid cultures (Figure 4J).

### Single-cell RNAseq analysis revealed higher metabolic dependency of AAM cells associated with FAO and NAD metabolism, which can be therapeutically targeted

Next, we evaluated the transcriptional differences between AAM and monocytes isolated from healthy donors (HD), using previously published single-cell RNA sequencing datasets (37). Unsupervised cluster analysis allowed the identification of 18 clusters, of which 6 were composed of cells enriched for macrophage/monocyte signatures (data not shown). Re-analysis of these within the 6 clusters (5777 AAMs and 814 healthy BM macrophages) using the Seurat analysis resulted in the identification of 12 new clusters (Figure 5A-B). Overall, 5 clusters exhibited the presence of HD macrophages as well as AAMs, whereas 7 clusters where mainly formed by AAMs (Figure 5C). Focusing on the clusters that contained a significant fraction of HD macrophages, differential gene expression analysis revealed upregulation of a limited number of genes in AAMs (Figure 5D). Among the up-regulated genes in AAMs we identified the M2 markers *CD163* and *MRC1* (*CD206*), genes associated with mitochondrial transfer (*EXOC5*), fatty acid metabolism (*CD36*), and the NAD^+^ salvage pathway gene nicotinamide phosphoribosyltransferase (*NAMPT*) (Figure 5E). Single cell gene set enrichment analysis (scGSEA) revealed enrichment for M1 macrophage and glycolytic metabolism signatures in HD macrophages and M2 macrophage and fatty acid metabolism signatures in AAM (Figure 5F-G), suggesting distinct polarization and metabolic profiles between AAMs and HD macrophages. Among the identified differentially expressed genes, NAMPT inhibition was recently identified as a metabolic vulnerability in LSCs by disrupting lipid homeostasis (38). Since the NAMPT inhibitor KPT-9274 showed promising results regarding apoptosis induction in LSC, sparing the heathy HSCs, we decided to evaluate the effects of KPT-9274 on AAMs as well. *Ex vivo* treatment of 25 AML patient samples with different chemotherapeutic and metabolism related drugs, revealed that patients with a high percentage of AAM exhibited increased resistance to Cytarabine (AraC) and venetoclax (VEN), and increased sensitivity to some macrophage targeting drugs (39) and KPT-9274 (Figure 5H). FACS analysis revealed that while blasts cells where equally sensitive to AraC, VEN, and KPT-9274, the AAM population was only targeted by KPT-9724, with no effects for AraC and VEN (Figure 5I). Although the treatment with KPT-9724 resulted in minimal effects on the surface marker expression profile of *in vitro* generated macrophages (Figure 5J), treated macrophages exhibited decreased supportive functions regarding cell proliferation and survival of AML cell lines, when compared to vehicle control (Figure 5K-L). Altogether, these results suggests that disruption of NAD^+^ generation can represent a therapeutical vulnerability via targeting of the supportive TME populations, providing new avenues for treatment.

**Figure 5.**
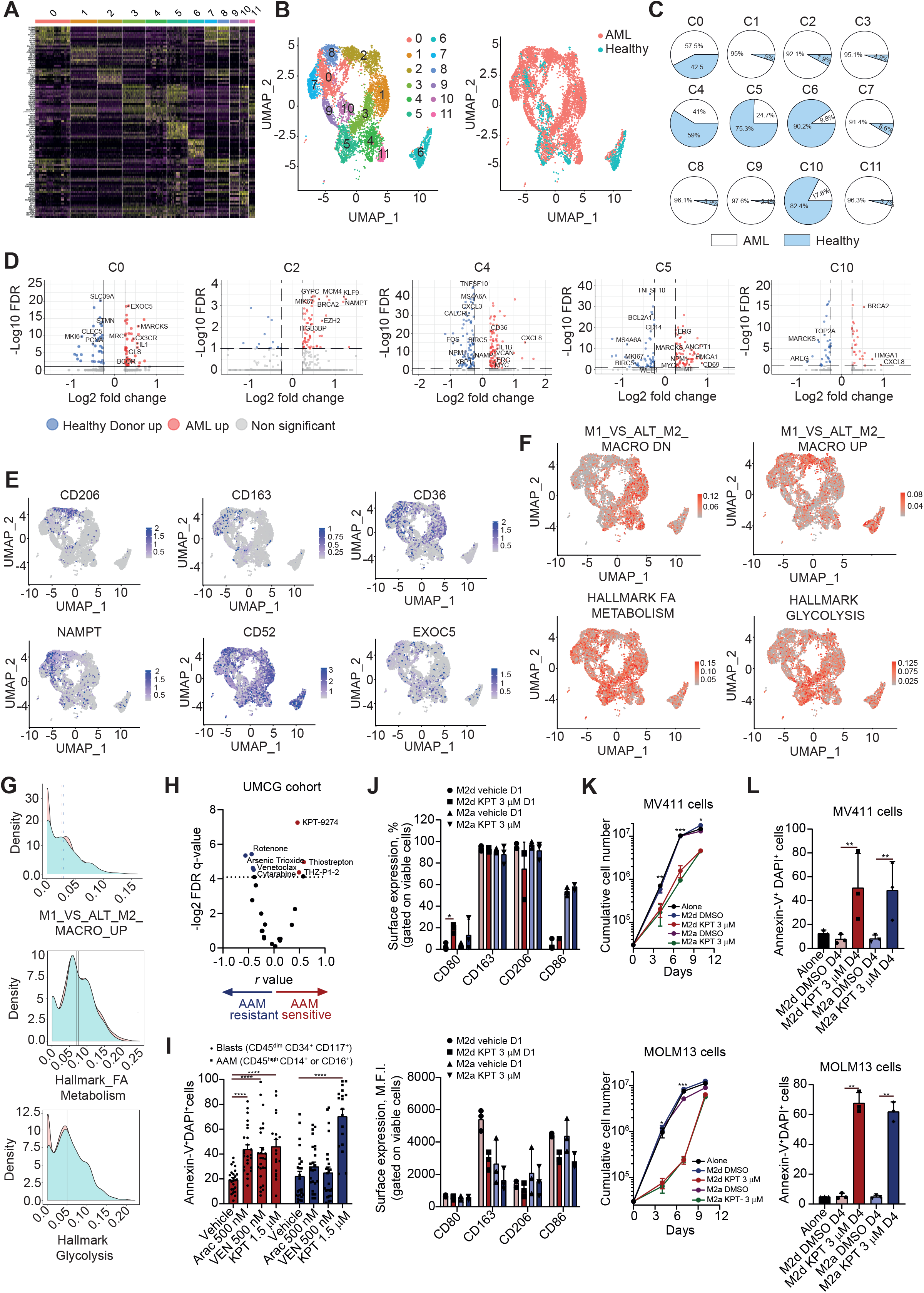
Single cell transcriptome analysis of AAMs and healthy macrophages. (A) Heatmap displaying the differentially expressed genes employed to perform the SEURAT clustering. (B) UMAP projection of monocytic/macrophage-like BM cells from 11 AML patients and 2 healthy donors, showing the formation of 12 main clusters. (C) Pie charts representing the frequency distribution of cells in each cluster, regarding the sample origin (Healthy donors vs AML). (D) Volcano plot demonstrating the differentially expressed genes in monocytes/macrophages from AML patients compared to healthy BM donors. (E) UMAP projection of differentially expressed genes in AAMs. (F) Single cell gene set enrichment analysis (scGSEA) projection in the macrophage landscape of AML patients and Healthy donors. (G) Density plots displaying the enrichment scores for the scGSEA. (H) Volcano plot demonstrating the Pearson correlation between the M2-macrophage levels (% by FACS) and the *ex-vivo* drug induced apoptosis in our set of primary AML samples (n=25 samples). Blue dots indicate significant negative correlation and red dot indicate a positive correlation. (I) Apoptosis (defined by Annexin-V^+^ alone or in combination with DAPI^+^) induced in AML leukemic blasts but not in the tumor associated macrophages (AAMs) after ex-vivo treatment with Cytarabine (Ara-C, dose: 500 nM) and Venetoclax (VEN, dose: 500 nM), and KPT-9274 (1.5 μM) for 72 hours (n = 25 samples). (J) Bar plot comparing the levels (%, upper panel; MFI, lower panel) of M2- (CD163 and CD206) and M1- (CD80 and CD86) markers measured by FACS in healthy activated M2a- and M2d-macrophages treated with vehicle and KPT-9274 (3 μM) for 24 h. (K) Cumulative cell count of MV4-11 (upper panel) and MOLM13 (lower panel) AML cell lines cultured on M2a and M2d macrophages treated with KPT-9274 (3 μM) for 24 h or vehicle (DMSO), prior to co-culture or alone (control) for 10 days. (L) Percentage of apoptosis induction on MV4-11 (upper panel) and MOLM13 (lower panel) cells after 4 days of culture in M2a and M2d macrophages treated with KPT-9274 (3 μM) for 24 h or vehicle (DMSO), prior to co-culture or alone (control) for 4 days.

Among the AAM correlated GSEAs, fatty acid oxidation (FAO) is one of the main mechanisms used by macrophages to fuel the tricarboxylic acid cycle, culminating in M2 polarization (40). To evaluate whether FAO is the driver of OXPHOS in M2d-exposed AML cells we treated M2d macrophages with the FAO inhibitor etomoxir (Eto) for 24h. After 24h macrophages were washed and co-cultured with primary AML cells for 24h to measure their functional respiration. Etomoxir-treated M2d macrophages exhibited decreased OCR and a decrease of M2 surface marker expression (supplemental Figure 5A-B). Also, OCR was not increased in HL60 or primary AML cells co-cultured on Etomoxir-treated macrophages (supplemental Figure 5C). Finally, the pre-treatment of M2d macrophages with Etomoxir abrogated the supportive properties to co-cultured AML blasts in long-term proliferation assays (supplemental Figure 5D).

### Clinical application of an M2 macrophage signature

To reduce the CIBERSORT M2 macrophage signature defined by over 500 genes, we aimed to generate a new simplified and clinically applicable M2 signature by selecting genes exclusively expressed by M2 macrophages (92 genes) from four different datasets (FANTON, BLUEPRINT, CIBERSORT and HPCA) (Figure 6A). From these 92, four genes (*CD52, FGR, GASK1B* and *RASA3*) could predict OS in at least 2 AML cohorts and were differentially expressed between the paired CD34^+^ or CD34^-^ compartments of AML patients (Figure 6B-C). We next used the combined gene expression of these 4 AAM-associated genes to design a prognostic score (M2-UMCG signature) to stratify AML patients regarding the clinical outcomes. The new M2-UMCG signature could independently predict OS and DFS survival in a “training” cohort (TCGA), considering ELN-risk stratification, age and sex as confounders (Figure 6D-E). Moreover, ROC curve analysis indicated superior OS prediction compared to the M2 CIBERSORT signature and the ELN2010 risk stratification (Figure 6F). We also confirmed that our signature is specific for M2 macrophages (supplemental Figure 6A) and validated our results in the BeatAML cohort (supplemental Figure 6B-C). Importantly, except of *GASK1B*, all other genes encode for plasma membrane proteins, and detection can easily be implemented in routine diagnostics. Additionally, since a growing body of evidence suggests that the remodeling of the BM niche impacts on the transition from myelodysplastic syndrome (MDS) to AML, we tested our M2-UMCG signature in MDS patients. Indeed, the M2-UMCG signature can also predict OS and progression-free survival (PFS) in MDS patients (Figure 6G). These results were specific to the M2 signature and independent from the revised International Prognostic Scoring System (IPSS-R)(supplemental Figure 6D-E). Our results show that incorporation of the TME as a clinical variable can improve AML and MDS risk stratification, facilitate the design of personalized treatment regiments, and provide the possibility of drug repurposing.

**Figure 6.**
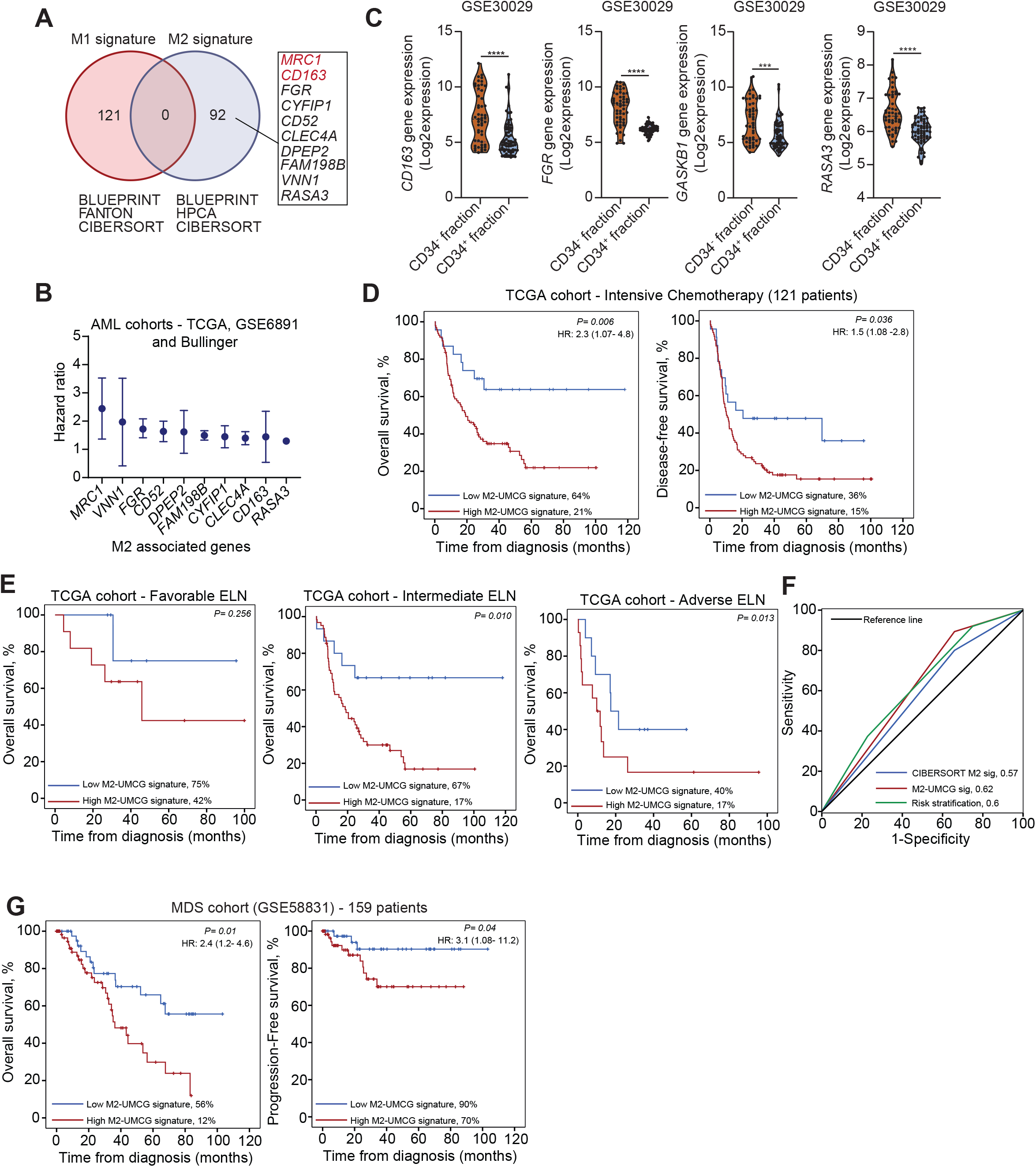
Development of an M2 signature for AML patients. (A) Venn diagram of genes exclusively expressed by M1 and M2 macrophages using the BLUEPRINT, FANTON, HPCA and CIBERSORT datasets. (B) Forest plot depicting the HR of each M2-associated gene (n=10) able to predict OS in at least 2 AML cohorts. (C) Expression of M2 genes in the CD34^-^/CD34^+^ compartments of the GSE30029 transcriptome set in which AML patient sample were sorted into paired CD34^+^ and CD34^-^ fractions. (D) KM analysis of OS and DSF using the M2-UMCG signature in the TCGA cohort. (E) KM analysis of OS within different ELN risk groups (favorable, intermediate, and adverse) using the M2-UMCG signature in the TCGA cohort. (F) Area under receiver operating characteristic (AUROC) curve plot of the predictive capacity of the M2-UMCG signature (red curve) *versus* the CIBERSORT M2 signature (blue curve) and the ELN2010 risk stratification (green curve) for OS in the TCGA cohort. AUROC=1 denotes perfect prediction and AUROC=0.5 denotes no predictive ability. (G) KM analysis of OS and progression-free survival using the M2-UMCG signature in the MDS cohort (GSE58831). (C) Mann-Whitney test. \*\**P* <0.01, \*\*\**P* <0.0001, NS, not significant. (D-E and G) Patients were dichotomized into low and high M2-UMCG signature. Survival curves were estimated using the KM method, and the log-rank test was used for comparison.

## Discussion

Here we show that macrophages strongly impact on the fate of LSCs by enhancing stemness, homing, immune evasion capacities and their metabolism. *In vivo* intra-BM co-injection of M2d-macrophages allowed the induction of full-blown leukemia, while APL blasts without co-injected macrophages were unable to expand in NSGS mice. The leukemic burden was consistently superior in M2d injected bones compared to M0-bones, suggesting that in particular M2d-type macrophages provide the leukemia-propagating signals. Perhaps even more intriguingly, even a two-day *in vitro* exposure of APL and AML blasts to M2d-macrophages - which we refer to as “training” - allowed efficient engraftment followed by fatal leukemia in NSGS mice. Apparently, the fate of leukemic blasts when encountering M2-macrophages can be two-fold: either they are phagocytosed, or they are altered in such a way that their leukemic potential has increased. In line with the substantial genetic heterogeneity among leukemia patients, we also identify clear heterogeneity in the macrophage landscape across different AML patients, whereby those with the highest proportion of tumor supportive M2 macrophages also display the poorest prognosis.

Our study might seem contradictory at first compared to the recent study published by Moore et al., showing improved engraftment of murine AML cells when macrophages are depleted in C57/BL6 mice (33). The authors demonstrate that inhibition of LC3-associated macrophage mediated phagocytosis resulted in the accumulation of apoptotic cells/bodies, which in return activated the STING pathway and promoted leukemia progression. While our data primarily focusses on human AML samples and xenograft models, we also observe a drastic decrease of AML cells when exposed to macrophages *in vitro* due to phagocytosis. Yet, the remaining cells are either a selection of leukemia inducing cells or have been transformed as such. The former would suggest that the cells were already present in the bulk and yet our control mice did not develop leukemia reinforcing the latter hypothesis. Thus, while the presence of macrophages can contribute to the reduction of tumor burden initially, the establishment of a leukemia supportive niche might be a matter of time and the long-term effects of M2 macrophages on AML cells *in vivo*, as our data suggests, could have detrimental effects on AML progression.

Chao et al., showed that the process of programmed cell removal relies on an equilibrium of pro and anti-phagocytic signals (34). While we did not observe an increase of “don’t eat me” signals, CALR expression was decreased after the two-day “training”, which could be attributed to an up-regulation of *STC1* (35). Furthermore, also α-integrins have been identified to regulate the expression of surface CALR in T-lymphoblasts during immunogenic cell death, whereby CALR binds to the α-integrin GFFKR motif preventing translocation of CALR to the surface (41). We show that M2d macrophages can upregulate the protein expression of CD49d-f on AML cells, which could be partly responsible for the reduction of surface CALR. More recently, Wattrus et al. identified CALR expressed by HSC to interact with the Lrp1ab present on macrophages in a brainbow zebrafish model. HSCs were either completely engulfed or a portion of the HSC was removed to promote HSC cell cycle progression (42). It will be of interest to further functionally study the role of CALR-mediated interactions between leukemic blasts and macrophages in future studies.

Although poorly understood, it has been noted that malignant tumor cells can fuse with healthy somatic cells to generate a more aggressive hybrid tumor cell. Here, we show that, like MSCs (36), macrophages can transfer mitochondria to AML blast cells driving OXPHOS metabolism in AML cells. In a study published by Tscheng et al, the authors validate the importance of very long chain acyl–CoA dehydrogenase (VLCAD) to promote AML proliferation, clonogenic growth and engraftment (43). Our data indicate that the two-day co-culture of primary AML cells on M2d macrophages is sufficient to induce long-term effects on AML proliferation, which could be abrogated by the pretreatment of macrophages with KPT-9274 and Etomoxir, thereby inhibiting OXPHOS. Overall, it is conceivable that the metabolic changes increase the proliferation capacity of leukemic cells resulting in a more aggressive leukemia. Of note, our single-cell analysis focusing on the AAMs, revealed that these cells rely on FAO and NAD^+^ generation to support their metabolic needs. Therefore, therapeutical strategies targeting those pathways could be clinically interesting for AML patients with high M2-profiles, representing a dual strategy of targeting the tumor cells and the TME as well, reducing the chances of a future niche supported relapse event. Together, these data indicate that direct interactions with macrophages can have a profound impact on tumor cell biology, in part mediated via the exchange of organelles such as mitochondria. These data corroborate with a previous study, which demonstrated that the co-culture of human CB CD34^+^ on M2 macrophages significantly increased the number of CD34^+^ and long-term HSCs (44). On the other hand, it is also quite conceivable that exchange of other factors, including plasma membrane proteins, participate in the aggressive leukemic phenotype that cells adopt after exposure to M2 macrophages, which will be investigated in future studies.

Congruent with the notion that AML is a highly heterogeneous disease we could classify patients based on their macrophage marker expression into the following groups: M2^high^M1^low^, M2^high^M1^high^ or M2^low^M1^high^. Within this context, scRNA sequencing analysis of AML patients identified 10 different subsets of macrophages with M2-like and immunosuppressive characteristics of which four could be identified by our FACS panel (CD163^high^, CD206^+^, CD14^high^ and CD16^+^) (17). A detailed functional understanding of these macrophage subtypes would be of interest to effectively target these AML supportive subpopulations. Here, we chose to first understand how healthy macrophages can affect the biology of AML blast cells assuming that leukemic cells are initially surrounded by healthy macrophages. Due to the inherent plasticity of macrophages, one could imagine that the malignant transformation generates AML supportive macrophages which in response promote the expansion of leukemic cells. Nevertheless, it remains to be determined whether the macrophages detected at diagnosis are wild-type (WT) or an intrinsic part of the leukemic clone. While leukemias are characterized by impaired differentiation, it has been described that some cells can escape the differentiation block (45). A study published by Van Galen et al., isolated CD14^-^ and CD14^+^ from AML patients and demonstrated that solely the CD14^+^ cells were able to inhibit T-cell activation suggesting immune modulatory functions of these cells. AML CD14^+^ monocytes showed transcriptional similarities with healthy monocytes with the exception of cytotoxic gene signatures being downregulated (37). Hence, it is conceivable that the leukemic cells can generate a tumor supportive microenvironment and raises the question whether malignant macrophages can provide even better tumor support than WT M2-like macrophages.

Finally, with immunotherapies being on the rise, the identification of clinical predictive markers becomes indispensable. Currently, the AML risk stratification system relies on genetic alterations detected in the leukemic blast cell itself to infer the course of disease progression (1). Here, we generated an M2-UMCG-signature to evaluate the prognostic value of the TME and correlated our signature to the therapeutic efficacy of several drugs. Our results show that incorporation of the TME as clinical variable can improve AML risk stratification, facilitate the design of personalized treatment regiments, and provide the possibility of drug repurposing.

## Methods

### Cell lines

All cell cultures were maintained in a humidified atmosphere at 37°C with 5% CO2. Mycoplasma contamination was routinely tested. All leukemia cell lines were authenticated by short tandem repeat analysis. The HS27A (CRL-2496) and HL60 (CCL-240™) cell lines were obtained from the American Type Culture Collection and grown in DMEM (for HS27A; Gibco, USA) or RPMI (for HL60; Gibco, USA) with 10% FBS. The MOLM13 (ACC 554) cell line was obtained from the DSMZ-German Collection of Microorganisms and Cell Cultures. *All-trans* retinoic acid (ATRA), Midostaurin (PKC), and arsenic trioxide (ATO) were obtained from Sigma-Aldrich (St. Louis, USA) and cytarabine (citarax) was obtained from Blau pharmaceuticals (Sao Paulo, Brazil). Venetoclax (VEN) and quizartinib (AC220) were obtained from Selleckchem (Houston, USA). Etomoxir was obtained from MedChemExpress (Groningen, NL).

### Human macrophage generation

Mononuclear cells were isolated by a density gradient using Ficoll (Sigma-Aldrich) from healthy donors or allogeneic donors. Next, 2.5×10^6^ or 5×10^6^ mononuclear cells were seeded into 12 or 6-well plates and incubated for 3h at 37°C in RPMI medium supplemented with 10% FBS, 10% heat inactivated and filtered human serum (AB serum, Thermofisher) and 1% penicillin and streptomycin (PS). After 2h of incubation, the non-adherent cell fraction was removed, and the new RPMI medium supplemented with 10% FBS, 10% human serum and 1% penicillin and streptomycin was added. Additionally, 50 ng/mL of GM-CSF (Prepotech) or M-CSF (Prepotech/Immunotools) growth factors were added to the medium to generate pre-orientated M1 and M2 macrophages, respectively. Monocytes were differentiated into macrophages over a time span of 6 days and at day 3 half of the medium was renewed.

### Flow cytometry

Cryopreserved MNC fractions of AML/APL patients were thawed, resuspended in newborn calf serum (NCS) supplemented with DNase I (20 Units/mL), 4 μM MgSO_4_ and heparin (5 Units/mL) and incubated at 37°C for 15 minutes (min). To analyze the myeloid fraction of the AML bulk sample, 5×10^5^ mononuclear cells were blocked with human FcR blocking reagent (Miltenyi Biotec) for 5 min and stained in FACS buffer (Phosphate buffer saline - PBS plus 2 mM EDTA, 2% BSA, 0.02% NaN3) with the following antibodies: CD45-FITC, HLA-DR-PE, CD14-PerCP, CD16-APC-Cy7, CD163-PE-Cy7, CD206-BV421 and CD80-APC for 20 min at 4°C. A more mature myeloid population was detected based on the CD45 staining and inside this gate HLA-DR positive cells were selected to analyze the CD14 *versus* CD16 cellular distribution. The different CD14-CD16 populations were then analyzed for the M1 marker CD80 and the M2 markers CD163 and CD206 (Figure 1B). Fluorescence was measured on the BD LSRII or FACS CantoII and analyzed using Flow Jo (Tree Star, Inc). For each sample a minimum of 5000 events were acquired inside the SSC-A^high^ CD45^high^ HLA-DR^+^ population.

### Human macrophage generation

Mononuclear cells were isolated by a density gradient using Ficoll (Sigma-Aldrich) from healthy donors or allogeneic donors. Next, 2.5×10^6^ or 5×10^6^ mononuclear cells were seeded into 12 or 6-well plates and incubated for 3h at 37°C in RPMI medium supplemented with 10% FBS, 10% heat inactivated and filtered human serum (AB serum, Thermofisher) and 1% penicillin and streptomycin (PS). After 2h of incubation, the non-adherent cell fraction was removed, and the new RPMI medium supplemented with 10% FBS, 10% human serum and 1% penicillin and streptomycin was added. Additionally, 50 ng/mL of GM-CSF (Prepotech) or M-CSF (Prepotech/Immunotools) growth factors were added to the medium to generate pre-orientated M1 and M2 macrophages, respectively. Monocytes were differentiated into macrophages over a time span of 6 days and at day 3 half of the medium was renewed.

### Murine macrophage generation

Mice were anesthetized with an overdose of ketamine/xylazine solution and euthanatized by cervical dislocation to collect femur and tibia. Next the epiphyses were cut off and a syringe of 25 G filled with PBS (1% FBS) was used to flush the BM onto a 70 μM cell strainer placed on a 50 ml tube. Red blood cells were lysed for 10 min at 4°C and washed with PBS. Three million BM mononuclear cells were seeded in a 100×20 mm petri dish and cultured in RPMI medium supplemented with 10% FBS, 15% of L929 supernatant and 1% PS for 7 days. At day 3 half of the medium was renewed.

### Macrophage polarization

At day 6 human macrophages were washed with PBS and new RPMI medium supplemented with 10% FBS was added. GM-CSF cultured macrophages were polarized to M1 macrophages with 20 ng/ml IFN-y (Prepotech) and 100 ng/ml LPS (Sigma-Aldrich), while M-CSF cultured macrophages were polarized to M2d macrophages with 10 ng/ml M-CSF and 20 ng/ml IL-6 (Prepotech). To generate M0 macrophages, M-CSF cultured macrophages were kept with 50 ng/mL of M-CSF in the medium.

Likewise, murine macrophages were washed with PBS and new RPMI medium supplemented with 10% FBS and 1% PS was added at day 6. To polarize murine macrophages to M2d macrophages, 20 ng/mL Il-6 (Prepotech) was added to the medium. M0 macrophages were generated by adding 15% of L929 supernatant.

### *In vivo* intra-BM APL patient derived xenotransplant (PDX) model

Macrophages were then detached with TrypLE (ThermoFisher) and washed 2 times with PBS at 450g. Macrophages were stained with DAPI, HLA-DR and CD206 to confirm viability and the macrophage subtype. Macrophages were resuspended in PBS at a at a working concentration of 1×10^5^/10 μL. Eight to ten weeks old female NSGS (NOD.Cg-Prkdcscid Il2rgtm1Wjl Tg(CMV-IL3,CSF2,KITLG)1Eav/MloySzJ) mice were anesthetized and 25 mg/Kg of Tramadol was injected subcutaneously. The intra-tibial injection of macrophages was performed according to the percutaneous approach (46). In summary, under anesthesia through a nose cone the mouse was placed in a supine position and the pre-shaved knee was cleaned with 70% ethanol and maintained in a flexed position. A 30 G needle was placed percutaneously through the knee joint and inserted by rotating the syringe. Once the BM space was reached, 10 μL were of the BM were aspirated (to free BM space). Next, 10 μL of macrophages corresponding to 100 000 cells in total were injected into the BM with a new needle. M0 macrophages were injected into the left tibia, while M2d macrophages were injected into the right tibia. Control mice received 10 μL of PBS. On the next day, six different cryopreserved MNC fractions of APL patient samples were thaw and depleted for CD3^+^ cells. Next, 1×10^6^ primary APL cells were injected via the retro-orbital sinus into macrophage recipient and control mice. Human CD45^+^ levels were measured regularly in peripheral blood samples obtained by sub-mandibular bleeding. Twelve weeks post-transplant mice were euthanized, and each leg of the macrophage recipient mice was processed separately. Cells from BM, spleen, and spine were collected and stained for human CD45, CD11b, HLA-DR, CD33 and CD117 to detect human APL blast cells (CD45^+^CD33^+^CD117^+^CD11b^-^HLA-DR^-^ cells). All specimens were acquired by flow cytometry (FACS CantoII) and analysed with the FlowJo software (Treestar, Inc., USA). For each sample a minimum of 20 000 viable events were acquired. In addition, cytospin preparations stained with May-Grünwald-Giemsa (MGG) were used to evaluate the morphology of human APL blasts.

### *In vivo* pre-culture APL PDX model

Human macrophages were generated and polarized as described in the sections “Macrophage generation” and “Macrophage polarization” in 6-well plates. APL patient samples were thaw as described in the section “Flow cytometry” (clinical characteristics in Supplemental table 1) and depleted for CD3^+^ cells. In total 5×10^6^ APL cells were put in co-culture per well for 48 hours. The co-cultures were performed in Gartner’s medium. After 48 hours, suspension cells were collected and washed 2x with PBS. Primary APL blast cells were counted and 1×10^5^ APL cells exposed to macrophages were set aside to evaluate the purity of these cells to ensure no macrophage contamination. Next, 1.5×10^5^ APL cells were injected via the retro-orbital sinus into 8 to 10 weeks old female NSGS (NOD.Cg-Prkdcscid Il2rgtm1Wjl Tg(CMV-IL3,CSF2,KITLG)1Eav/MloySzJ) mice. For control mice the respective paired APL sample, which was used for co-culture was thaw as described in the section “Flow cytometry” and depleted for CD3^+^ cells. A total of 1×10^6^ cells were transplanted via the retro-orbital sinus directly after thawing and the rest of the cells were put in co-culture with primary human mesenchymal stromal cells for 48 hours. The same culture conditions were applied as for the macrophage co-culture. After 48 hours cells were collected and counted to inject 200-350 000 APL blast cells via the retro-orbital sinus. Monitoring of APL engraftment and evaluation of APL blast infiltration post-sacrifice was executed as described in the section “In vivo intra-BM APL PDX model”.

### *In vivo* Limiting dilution assay

Engrafted primary APL blast cells, which were pre-cultured on M2d macrophages and induced fatal leukemia were sorted post-sacrifice based on the human markers CD45, CD33 and CD117. Sorted APL blast cells from primary transplant were then transplanted via the retro-orbital sinus in secondary mice at different cell dosage: 1×10^3^, 1×10^5^ and 1.5×10^6^. Control mice received different cell dosage as well: 5×10^5^, 1×10^6^ and 5×10^6^ of APL samples at diagnosis. The frequency of leukemic initiating stem cells was calculated with the ELDA software (47).

### Oxygen consumption (OCR) and extracellular acidification rate (ECAR) measurements

Oxygen consumption rate (OCR) and Extra Cellular Acidification Rate (ECAR) were measured using Seahorse XF96 analyzer (Seahorse Bioscience, Agilent, US) at 37 °C. For AML cell lines (CD45^+^HLA-DR^-^ cells) and sorted CD34^+^ or CD117^+^ from primary AML patients, 1×10^5^ and 1.5×10^5^ viable cells (DAPI^-^) were seeded per well in poly-L-lysine (Sigma-Aldrich) coated Seahorse XF96 plates in 200 μL XF Assay Medium (Modified DMEM, Seahorse Bioscience), respectively. For OCR measurements, XF Assay Medium was supplemented with 10 mM Glucose and 2.5 μM oligomycin A (Port A), 2.5 μM FCCP (carbonyl cyanide-4-(trifluorometh oxy) phenylhydrazone) (Port B) and 2 μM antimycin A together with 2 μM Rotenone (Port C) were sequentially injected in 25 μL volume to measure basal and maximal OCR levels (all reagents from Sigma-Aldrich). For ECAR measurements, Glucose-free XF Assay medium was added to the cells and 10 mM Glucose (Port A), 2.5 μM oligomycin A (Port B) and 100 mM 2-deoxy-D-glucose (Port C) were sequentially injected in 25 μL volume (all reagents from Sigma-Aldrich). For measuring the metabolic activity after M2d or MS-5 co-culture, we initially seeded 2.5×10^5^ – 5×10^5^ AML cells (cell lines and primary samples, respectively) for 2 days and counted remaining viable cells, loading equal amounts of tumor cells. For etomoxir experiments, MS5 and M2d macrophages were treated with Etomoxir (50 μM) alone for 24 h. After that, cells were washed twice with 1X PBS and HL60 cells or primary AML blasts were added and a co-culture for 24 h was performed, to measure the OCR/ECAR in the AML cells. To exclude off-target effects from treated MS5 or M2d macrophages on the exposed AML cells after coculture, AML cells were evaluated with the mitochondrial markers – MitoTracker DeepRed and Tetramethylrhodamine, Ethyl Ester, Perchlorate (TMRE; ThermoFisher) by FACS (LSRII). All XF96 protocols consisted of 4 times mix (2 min) and measurement (2 min) cycles, allowing for determination of OCR at basal and also in between injections. Both basal and maximal OCR levels were calculated by assessing metabolic response of the cells in accordance with the manufacturer’s suggestions. The OCR measurements were normalized to the viable number of cells used for the assay.

### Mitochondrial transfer assay

Mitochondrial transfer assays were performed as described by Moschoi et al (36). Co-cultures of primary AML blasts or cell lines with M2d macrophages and MS-5 confluent monolayer were performed in Gartners medium or RPMI 1640, respectively. Etomoxir (50 μM; MedChemExpress, NL) treatment were performed in the MS5 and M2d macrophages 24h prior to the co-culture. Viability of the stromal cells was evaluated by DAPI staining. MS-5 and M2d macrophage MitoTracker loading was performed as follows: confluent stromal cells were stained for 10 minutes with 2 μM MitoTracker Green FM and 1 μM MitoTracker DeepRed FM (Molecular Probes), washed twice, and left 72 hours to allow elimination of the unbound probe. Stromal cells were then washed twice again before initiating co-cultures with AML cells. As a quality control, conditioned medium of stained MS-5 and M2d macrophages 72 hours post staining, was collected and used to stain AML cells, to evaluate the leakage of MitoTracker dyes.

### *In vivo* homing assay

In total 5×10^6^ CD3^+^ depleted APL cells were put in co-culture on M2d macrophages per well for 48 hours. After 48 hours, suspension APL cells were collected and counted. A total of 1×10^6^ co-cultured APL blast cells were washed 2x in serum free medium at 450g for 5 min. APL cells were resuspended in 100 μL serum free medium and stained with 1 μL Incucyte® Cytolight Rapid Red Dye (0.33 μM) for 20 min at 37°C. Cells were then washed 2x in medium with serum at 450g for 5 min. For the control group cryopreserved MNC fractions of APL patient samples were thaw, depleted for CD3^+^ and 1×10^6^ were labelled with the Incucyte dye as described above. Labeled APL blast cells (after thawing and M2d co-cultured) were transplanted via the retro-orbital sinus. Additional, 0.25-1×10^6^ freshly thaw and co-cultured blast cells were set aside to stain for purity and measure the levels of CD49d, CD49e and CD49f. All specimens were acquired by flow cytometry (FACS CantoII) and analysed with the FlowJo software (Treestar, Inc., USA). For each sample a minimum of 20 000 events were acquired. Eighteen hours post-transplant mice were sacrificed, and legs were first flushed and then crushed to retrieve the maximum number of cells. BM cells were stained with DAPI and human CD45-APC, CD33-PE and the Incucyte dye. A total of 5×10^6^ cells DAPI negative cells were acquired by flow cytometry (FACS CantoII) and analysed with the FlowJo software (Treestar, Inc., USA).

### Immunohistochemistry

Tissue sections were cut from formalin fixed embedded BM biopsies of AML patients at diagnosis. For CD68 and CD163 were stain visualized by the Ventana Benchmark Ultra automated slide stainer, after antigen retrieval (Ultra CC1, Ventana Medical Systems), using monoclonal antibodies CD68 (PG-M1, 1:100, Dako) and CD163 (MRQ-26, ready to use, Ventana) and Ultraview (Ventana). Digital images of these slides were scored for percentage of positively staining BM cells.

### Statistics

For statistical analysis of 2 groups, either a paired Wilcoxon matched-pairs signed rank test, or an unpaired Mann-Whitney U test was used. When more than 2 groups were compared, a Kruskal-Wallis test (1-way ANOVA) was performed followed by a Dunn’s post hoc test for significance using GraphPad Prism, version 8.1 for Windows 10 (GraphPad Software). For the correlation analysis, a pearson correlation was drawn. Differences among group means were considered significant when the P value was less than 0.05.

### Study approval

Bone marrow samples of APL patients used in for *in vivo* experiments were studied after informed consent and protocol approval by the Ethical Committee in accordance with the Declaration of Helsinki (registry #12920; process number #13496/2005; CAAE: 155.0.004.000-05 and CAAE: 819878.5.1001.5440). Mononuclear cells (MNCs) were isolated via Ficoll (Sigma-Aldrich) separation and cryopreserved. Peripheral blood (PB) and bone marrow (BM) samples of AML patients were studied after informed consent and protocol approval by the Medical Ethical committee of the UMCG in accordance with the Declaration of Helsinki. An overview of patient characteristics can be found in Table 1.

**Table 1.**
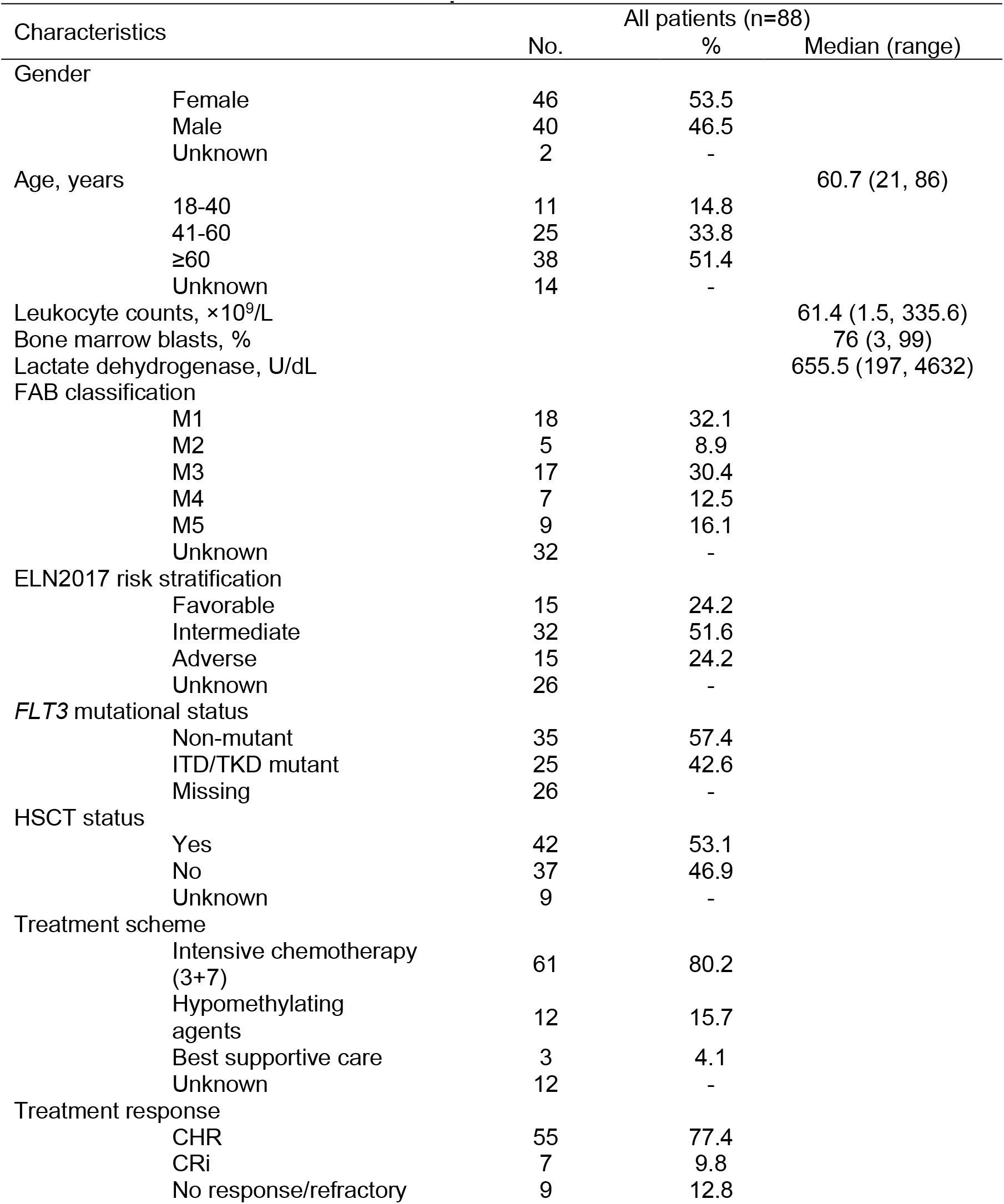
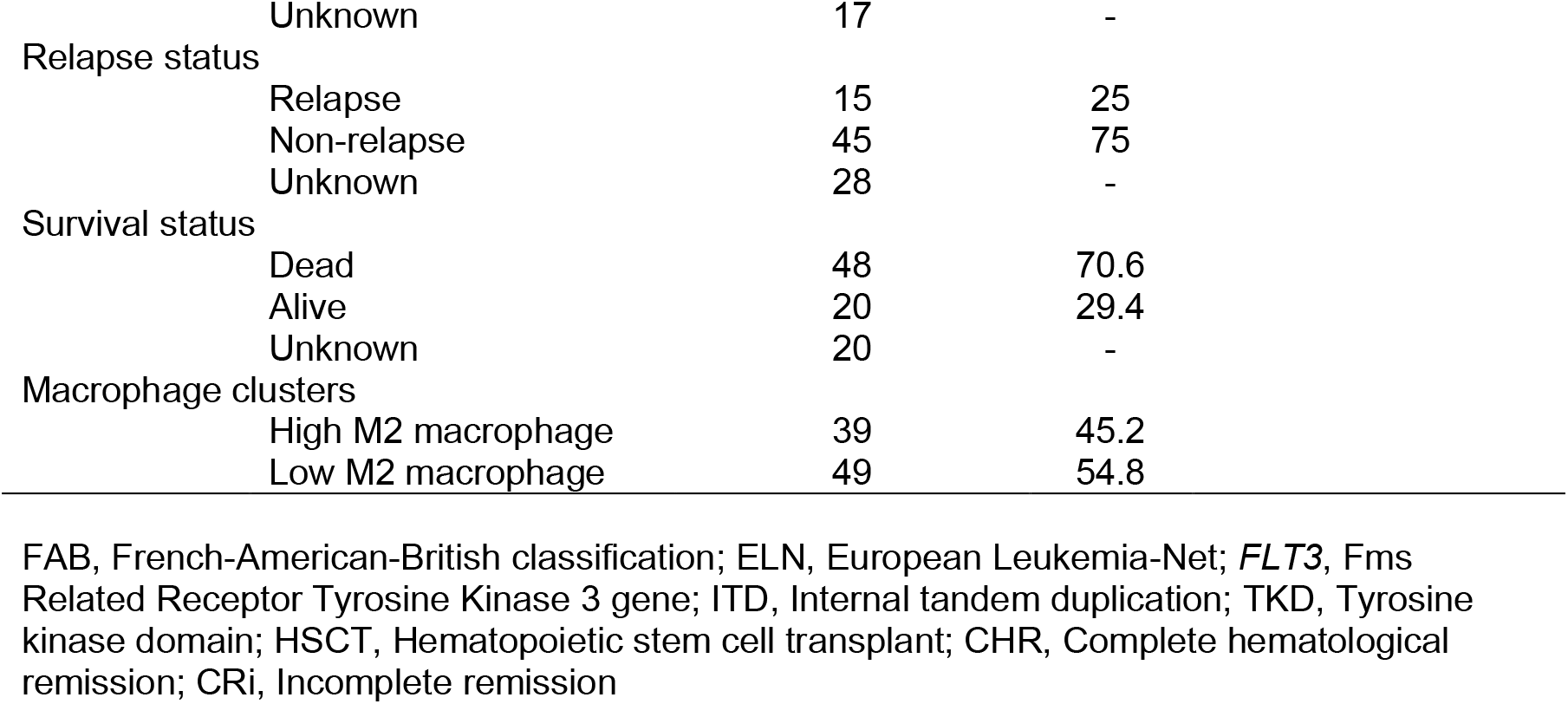
Clinical characteristics of AML patients included.

All animals were housed under specific pathogen free conditions in individually ventilated cages during the whole experiment. The animals were maintained according to the Guide for Care and Use of Laboratory Animals of the National Research Council, USA, and to the National Council of Animal Experiment Control recommendations. All experiments were approved by the Animal Ethics Committee of the University of São Paulo (protocols #176/2015 and #095/2018).

Additional methods can be found in the Supplemental materials and Methods

## Authorship contributions

I.W., D.A.P-M, E.M.R and J.J.S. conceived and designed the study, performed experiments, analyzed, and interpreted data, performed the statistical analyses, and drafted the article. L.Y.A, J.R.H, C.O, C.L.A, T.M.B, J.M.M, A.D, G.H, A.R.L-A, performed experiments, collected data, and reviewed the paper. D.R.S, L.Q, and S.M.H performed bioinformatics analysis and reviewed the paper. N.V. provided help with phagocytosis assays and reviewed the paper. A.D. performed immunohistochemistry analyses and reviewed the paper, J.R.H, E.A. and G.H. provided patient samples and clinical data and reviewed the paper. All authors gave final approval of the submitted manuscript.

## Acknowledgements

This investigation was supported by Fundação de Amparo à Pesquisa do Estado de São Paulo (FAPESP, Grant #2013/08135-2). I.W. received a fellowship from FAPESP (Grant #2015/09228-0). D.A.P-M. received a fellowship from FAPESP (Grant #2017/23117-1). I.W and D.A.P-M were sponsored by the Abel Tasman Talent Program (ATTP) of the Graduate School of Medical Sciences of the University of Groningen/University Medical Center Groningen (UG/UMCG), The Netherlands.

