## supplemental files for "M2-polarized macrophages control LSC fate by enhancing stemness, homing, immune evasion and metabolic reprogramming"

#### Resources table

| REAGENT or RESOURCE | SOURCE | IDENTIFIER |
|---|---|---|
| **Antibodies** | | |
| Anti-Human CD45 FITC (1:50 dilution) | BioLegend | 368508<br>RRID: AB_2566368 |
| Anti-Human CD45 PE-Cy7 (1:100 dilution) | BioLegend | 304016<br>RRID: AB_314404 |
| Anti-Human HLA-DR PE (1:50 dilution) | BioLegend | 980416<br>RRID: AB_2892517 |
| Anti-Human CD14 PerCP-Cy5 (1:50 dilution) | BioLegend | 301848<br>RRID: AB_2564059 |
| Anti-Human CD16 APC-Cy7 (1:50 dilution) | BioLegend | 302018<br>RRID: AB_314218 |
| Anti-Human CD163 PE-Cy7 (1:50 dilution) | BioLegend | 333614<br>RRID: AB_2562641 |
| Anti-Human CD206 BV421 (1:50 dilution) | BD Biosciences | 564062<br>RRID: AB_2738570 |
| Anti-Human CD80 APC (1:50 dilution) | BioLegend | 305220<br>RRID: AB_2076147 |
| Anti-Human CD47 PE (1:50 dilution) | BD Biosciences | 556046<br>RRID: AB_396317 |
| Anti-Human CD33 PE (1:20 dilution) | BD Biosciences | 561816<br>RRID:<br>AB_10896480 |
| Anti-Human CD117 APC (1:50 dilution) | BD Biosciences | 550412<br>RRID: AB_398461 |
| Anti-Human CD34 eFluor 450 (1:20 dilution) | eBiosciences | 48-0341-82<br>RRID: AB_2043837 |
| Anti-human CD11b FITC (1:20 dilution) | Immunotools | 21279113X2 |
| Anti-human CD11 Alexa Fluor 594 (1:50 dilution) | BioLegend | 101254<br>RRID: AB_2563231 |
| Annexin FITC (1:200) | Immunotools | 31490013X2 |
| Annexin APC (1:200) | Immunotools | 31490016X2 |
| Anti-human CD49d PE (1:50 dilution) | BioLegend | 304304<br>RRID: AB_314430 |
| Anti-human CD49e PE (1:50 dilution) | BioLegend | 328010<br>RRID: AB_2280539 |
| Anti-human CD49f PE (1:50 dilution) | BioLegend | 313612<br>RRID: AB_893373 |
| Anti-human CD24 PE (1:50 dilution) | BioLegend | 311106<br>RRID: AB_314855 |
| Anti-mouse Gr1 APC (1:100 dilution) | BioLegend | 108412<br>RRID: AB_313377 |
| Anti-mouse CD117 APC-Cy7 (1:100 dilution) | BioLegend | 105826<br>RRID: AB_1626278 |
| Anti-mouse CD16/32 FITC (1:100 dilution) | BioLegend | 101306<br>RRID: AB_312805 |
| Anti-mouse CD34 PE-Cy7 (1:100 dilution) | BioLegend | 128618<br>RRID: AB_2721678 |
| Anti-mouse CD45.1 PE (1:100 dilution) | BioLegend | 110708<br>RRID: AB_313497 |
| Anti-mouse CD45.2 FITC (1:100 dilution) | BioLegend | 109806<br>RRID: AB_313443 |
| Anti-human CD68 (IHC) (1:100 dilution) | DAKO | PG-M1<br>RRID: AB_2892683 |
| Anti-human CD163 (IHC) | Ventana | MRQ-26<br>RRID: AB_1159122 |

| | | |
|---|---|---|
| CALR Polyclonal Antibody (FACS/CM) | ThermoFisher | PA3-900<br>RRID: AB_325990 |
| Anti-Stanniocalcin 1 | Abcam | Ab83065<br>RRID: AB_1861344 |
| Donkey anti-Rabbit (H+L) AF647 | ThermoFisher | A32795<br>RRID: AB_2762835 |
| Rabbit anti-Mouse (H+L) AF594 | ThermoFisher | A27027<br>RRID: AB_2536090 |
| **Bacterial and virus strains** | | |
| **Biological samples** | | |
| Human AML blast cells | UMCG/USP | Ethical committee<br>NL43844.042.13 |
| Human APL blast cells | UMCG/USP | Ethical committee<br>#13496/2005 |
| Murine hCG-PML-RARa blast cells | USP | Ethical committee<br>#176/2015 |
| HSC from C57/BL6 mice | USP | Ethical committee<br>#176/2015 |
| **Chemicals, peptides, and recombinant proteins** | | |
| Propidium Iodide | ThermoFisher | P1304MP |
| 4′,6-diamidino-2-phenylindole | Sigma-Aldrich | 28718-90-3 |
| Paraformaldehyde | Sigma-Aldrich | 30525-89-4 |
| DNase I | Roche | 11284932001 |
| MgSO4 | Sigma-Aldrich | M7506 |
| Heparin | | |
| Arsenic Trioxide | Sigma-Aldrich | 1327-53-3 |
| All Trans Retinoic Acid | Sigma-Aldrich | 302-79-4 |
| Midostaurin | Sigma-Aldrich | M1323 |
| Quizartinib | Selleckchem | S1526 |
| Venetoclax | Selleckchem | S8048 |
| Etomoxir | MedChemExpress | HY-50200 |
| Tocilizumab (Actemra/RoActemra) | Roche | |
| Human Granulocyte-macrophage colony-stimulating factor | Peprotech | 300-03 |
| Human Macrophage Colony-Stimulating Factor | Immunotools | 11343117 |
| Human Interleukin 6 | Peprotech | 200-06 |
| Human Interferon Gamma | Peprotech | 300-02 |
| Human Interleukin 3 | Peprotech | 200-03 |
| Human Granulocyte colony-stimulating factor | Peprotech | 300-23 |
| Human Thrombopoietin | Amgen | |
| Lipopolysaccharides | Sigma-Aldrich | L4391 |
| Carboxyfluorescein succinimidyl ester | BioLegend | 423801 |
| Incucyte® Cytolight Rapid Red Dye | Sartorius | 4706 |
| Hydrocortisone | Sigma-Aldrich | 50-23-7 |
| β-mercaptoethanol | Merck Sharp & Dohme BV | 60-24-2 |
| SsoAdvanced Universal SYBR® Green Supermix | BioRad | 1725274 |
| iScript cDNA synthesis Kit | BioRad | 1708891BUN |
| Tetramethylrhodamine, Ethyl Ester, Perchlorate | Thermofisher | T669 |
| MitoTracker DeepRedTM | Thermofisher | M22426 |
| MitoTracker GreenTM | Thermofisher | M7514 |
| **Critical commercial assays** | | |

| | | |
|---|---|---|
| Seahorse XFe96 Flux Analyzer | Agilent | |
| **Deposited data** | | |
| RNA-seq data | Ley TJ et al., 2013 (PMID: 23634996) | https://tcga-data.nci.nih.gov/tcga/ |
| Single cell antibody sequencing data | Tirana et al., 2021 (PMID: 34811546) | https://abseqapp.shiny.embl.de/ |
| **Experimental models: Cell lines** | | |
| MOLM13 | DSMZ | ACC 554<br>RRID: CVCL_2119 |
| HL60 | ATCC | CCL-240™<br>RRID: CVCL_0002 |
| MS-5 | DSMZ | ACC 441<br>RRID: CVCL_2128 |
| HS27A | ATCC | CRL-9591™ |
| L929* | ATCC | CCL-1™ |
| HEK293T | ATCC | CRL-3216<br>RRID: CVCL_0063 |
| **Experimental models: Organisms/strains** | | |
| NSGS (NOD.Cg-Prkdcscid Il2rgtm1Wjl Tg(CMV-IL3,CSF2,KITLG)1Eav/MloySzJ) | Jackson Laboratory | 013062<br>RRID: IMSR_JAX:013062 |
| NSG (NOD.Cg-Prkdcscid Il2rgtm1Wjl/SzJ) | Jackson Laboratory | 005557<br>RRID: IMSR_JAX:005557 |
| C57BL/6J | Jackson Laboratory | 000664<br>RRID: IMSR_JAX:000664 |
| B6.SJL-Ptprca Pepcb/BoyJ | Jackson Laboratory | 002014<br>RRID: IMSR_JAX:002014 |
| **Oligonucleotides** | | |
| *STC1* Forward primer | Eurofins | TGCTAAATTTGACACTCAGGGAAA |
| *STC1* Reverse primer | Eurofins | ACCTCAGCAATCATCCTTTGG |
| *HPRT1* Forward primer | Eurofins | GAACGTCTTGCTCGAGATGTGA |
| *HPRT1* Reverse primer | Eurofins | TCCAGCAGGTCAGCAAAGAAT |
| *ACTB* Forward primer | Eurofins | AGGCCAACCGCAAGAAG |
| *ACTB* Reverse primer | Eurofins | ACAGCCTGGATAGCAACGTACA |
| *RPL30* Forward primer | Eurofins | ACTGCCCAGCTTTGAGGAAAT |
| *RPL30* Reverse primer | Eurofins | TGCCACTGTAGTGATGGACAC |
| **Recombinant DNA** | | |
| **Software and algorithms** | | |
| FlowJo v10.0.6 | Treestar | http://www.flowjo.com/ |
| Prism 9 | GraphPad | http://www.graphpad.com/ |
| SPSS Statistical package 19.1 | IBM | https://www.ibm.com/ |

| | | |
|---|---|---|
| Wave | Agilent | https://www.agilent.com/ |
| RStudio | CRAN | www.r-project.org |
| GSEA 4.0.1 | Broad Institute | https://software.broadinstitute.org/gsea/ |
| Cytoscape 3.4 | | http://apps.cytoscape.org/apps/bingo |
| **Other** | | |
| FcR Blocking reagent, human | Miltenyi Biotec | 130-059-901 RRID: AB_2892112 |
| CD34 MicroBeads Kit UltraPure, Human | Miltenyi Biotec | 130-100-453 |
| CD117 MicroBeads Kit, Human | Miltenyi Biotec | 130-091-332 |
| CD3 MicroBeads, Human | Miltenyi Biotec | 130-050-101 |
| Streptavidin MicroBeads | Miltenyi Biotec | 130-048-102 |
| TrypLE™ Express Enzyme | Thermo Fischer Scientific | 12604021 |
| MethoCult™ | Stemcell | H4435 |
| KAPA RNA HyperPrep Kit with RiboErase (HMR) | Roche | 08098131702 |

#### Resource availability

##### Lead contact

##### Materials availability

Materials generated in this study are available on request.

#### METHOD DETAILS

##### CIBERSORT analysis

Relative immune cell fractions were estimated using the CIBERSORT, quanTIseq, xCell, MCP-counter and EPIC algorithms, based on several reference expression signatures that distinguish at maximum 64 immune cell subtypes. Briefly, normalized gene expression data were uploaded to the TIMER2.0 web portal (http:// timer.cistrome.org/), with the data matrices prepared according to the example. After 1000 permutations, only samples with p-values < 0.05 were included in subsequent analyses. Kruskall-Wallis were applied to identify immune subpopulations that were differentially enriched between the different AML patients, controlling for the false discovery rate (FDR) by the Benjamini–Hochberg method (FDR < 0.05).

##### Murine *in vivo* pre-culture

Murine macrophages were generated and polarized in 100x20 mm petri dishes. Murine primary APL blasts were collected from hCG-PML-RARA mice (CD45.2 background), which developed APL. Once macrophages were polarized, the macrophages were washed and new RPMI supplemented with 10% FBS was added. Murine APL blast cells were then thaw and 5-10x$10^6$ were put in co-culture for 48h. After 48 hours, suspension cells were collected and washed 2x with PBS. Murine APL blast cells were counted and 1x$10^5$ APL cells exposed to macrophages were set aside to evaluate the purity of these cells. Next, 1.5x$10^5$ murine APL cells were injected via retro-orbital sinus into sub-lethally irradiated (350 cGy) C57BL/6J.PepBoy recipients (CD45.1). For control mice the respective paired murine APL sample, which was used for co-culture was thaw and a total of 1x$10^6$ cells were transplanted via the retro-orbital sinus directly after thawing. Chimerism levels were evaluated by measuring the CD45.2 marker by flow cytometry in blood samples obtained by sub-mandibular bleeding. Mice were followed for OS analysis and euthanized when tumor reached ethical limits (>90% engraftment) or mice showed severe signs of illness. At the end of the experiment, animals were harvested and the BM was analyzed for the presence of early and late promyelocytes defined by CD34 and CD16/32 ($CD34^+CD16/32^+$, Early Pro/$CD34^-CD16/32^+$, Late Pro) inside the population of lineage negative cells (lineage markers defined by CD3e, CD19, B220, Ter119, NK1.1, CD4, and CD8) positive for CD117 and Gr1 intermediate (*1, 2*).

**_In vitro_ phagocytosis**

Macrophages, which were generated in a 6 well plate were detached with TrypLE and $1x10^4$ macrophages were seeded in a flat-bottom 96 well plate to reach a confluence of approximately 80%. Macrophages were then incubated overnight to allow adherence. Cryopreserved MNC fractions of AML cells were thaw and $CD3^+$ depleted. For the phagocytosis at day 0, a total of $1x10^5$ AML cells were washed 2x in serum free medium at 450g for 5 min, resuspended in 100 µL and stained with 1 µL of Incucyte® Cytolight Rapid Red Dye (0.33 µM) or CSFE (1:100 dilution, ThermoFisher). Cells were incubated for 20 min at 37°C and washed 2x with medium supplemented with serum. A total of $5x10^4$ were co-cultured with macrophages in a 96-well plate and incubated for 3h at 37°C and 5% $CO_2$. After 3h of incubation AML cells were gently removed by washing with PBS 3x times. To visualize macrophages, residual cells were stained with CD11b-FITC (1 µg/mL) (when AML cells labeled with Incucyte red dye) or CD11b-AF594 (1 µg/mL) (when AML cells labeled with CSFE) 30 min at room temperature in the dark. After 30 min macrophages were fixed with 2% paraformaldehyde, resuspended in PBS with DAPI solution and three pictures of randomly chosen fields of view were taken by the EVOS Cell Imaging System (ThermoFisher). The percentage of phagocytosis was equal to the number of macrophages containing labeled AML cells per 100 macrophages. For phagocytosis at day 2, a total of $5x10^5$-$1x10^6$ AML cells were put in co-culture with either M2d macrophages or MS-5 cells seeded in a 12 well plate for 48h. The co-cultures were performed in Gartner's medium consisting of α-MEM (Thermo Scientific) supplemented with 12.5% fetal bovine serum (Gibco), 12.5% horse serum (Gibco), 1% penicillin and streptomycin, 2 mM glutamine (Gibco), 57.2 mM β-mercaptoethanol (Merck Sharp & Dohme BV), and 20 ng/mL G-SCF, TPO and IL-3.. After 48h of co-culture, AML cells were collected, and a phagocytosis assay was performed as described above. In addition to the phagocytosis assay, primary AML cells were stained for CD47 and CD24 at diagnosis and after two-day co-culture on M2d and MS5. All specimens were acquired by flow cytometry (BD LSR II) and analysed with the FlowJo software (Treestar, Inc., USA). For each sample a minimum of 10 000 events were acquired.

##### Colony forming unit assay

Primary $CD3^+$ depleted AML cells ($1x10^6$) were put in co-culture with either macrophages or MS5 cells for 48h. The co-cultures were performed in Gartner's medium. After 48h cells were collected, washed and counted. A total of $1x10^3$ AML cells post co-culture were plated in semisolid methylcellulose medium supplemented with human cytokines MethoCult$^{TM}$ H4435 (StemCell). Colonies were detected after 10 (For BFU-E colonies, when present) -14 days and scored.

##### *In vitro* AML proliferation in liquid culture

Primary $CD3^+$ depleted AML cells ($5x10^5$-$1x10^6$) were put in co-culture with either macrophages or MS5 cells for 48h. The co-cultures were performed in Gartner's medium. After 48h, cells were collected, washed and counted. A minimum of $1x10^5$ primary AML cells were put in liquid culture with Gartner's medium. Cultures were grown at 37°C and 5% $CO_2$ and demi-populated after counting if necessary. Cell proliferation was assessed with a hemocytometer until day 30.

##### ***In vitro*** **migration assay**

Primary BM stromal cells (passage 3) were plated at a density of $1x10^5$ per well in a transwell system in a 6-well plate format. Migration assay was performed as previously described(*3*). Briefly, primary AML/APL blasts were co-cultured with M2d macrophages for 48 h, and after the co-culture, diagnosis and co-cultured blasts were seeded ($1x10^6$ cells) in the upper chamber of the 6-well plate and incubated at 37˚C at 5% $CO_2$ for 18 hours to allow the migration upon the SDF-1 stimulus (produced by the MSCs in the lower chamber). After incubation, membranes were fixed with Methanol 99.9% and stained with Violet Crystal. Colonies containing at least 50 cells were counted and considered as migrating cells. The values were then normalized to the diagnosis samples.

##### **Development of a M2 signature suitable for AML patients**

The genetic signatures from M1 and M2 macrophages were retrieved from the FANTON, HPCA, BluePrint and CIBERSORT signatures(*4-6*). Genes with unique differential expression in M2 but not M1 macrophages, were selected for survival analysis using AML cohorts (Fig. 6a). Out of those genes, the *CD163*, *RASA3*, *FGR* and *GSK1B* (also called *FAM198B*) were able to predict poor OS in at least 2 independent AML cohorts and exhibited increased expression in the $CD34^-$ negative cells, when compared to the $CD34^+$ cells in AML bone marrows (Fig. 6b-c). We combined the gene expression of those four M2-associated genes to create a prognostic score (M2-UMCG signature) able to stratify AML patients regarding the clinical outcomes and M2-macrophage genetic signature profile. Next, using the TCGA cohort (*7*), patients were dichotomized as a low or high expression using receiving operating characteristics (ROC) curve and the C index and were interrogated for univariate and multivariate Cox proportional hazards model (CPHM) regression analyses for overall survival (OS). Including age, sex and European LeukemiaNet (ELN2010) as cofounders, we identified the independent prognostic predictors by backward elimination using an exclusion significance level of 5%. The M2-UMCG signature was defined as the weighted sums of hazard ratio from the final Cox model from independent prognostic genes. Internal validation was performed using a non-parametric bootstrap procedure with 1,000 resamplings to get estimates of HR between risk categories corrected for overfitting.

We used the BeatAML cohort (*8*) and the MDS cohort (GSE58831) to validate our M2-UMCG signature (Fig. 6f). For the BeatAML cohort, was considered the ELN2017 risk-stratification, age and sex as confounders in the multivariate CPHM. Moreover, we correlated the M2-UMCG signature with the drug sensitivity against the 124 compounds evaluated in the BeatAML study, to identify the possible therapeutic opportunities for AML patients with high M2 signature. Area under the curve values (AUC) were used to perform the Pearson correlation with the M2-UMCG signature.

#### Supplemental Figure Legends

**Fig. S1. Extended analyses of the macrophage landscape in AML.**

(A) Heat map showing the expression of CD163, CD206 and CD80 detected in monocytes/macrophages of H.D and AML patients. (B) M1 (upper panel) and M2 macrophage (lower panel) levels from primary AML samples (n=67 samples) were compared among a multitude of genetic alterations. Significance was determined using either two-tailed Mann–Whitney or Kruskal–Wallis tests (for categorical variables). Blue indicates ELN risk-stratification groups and red indicates the different genetic mutations evaluated. Only mutations that were found in ≥3 patients were considered. Significance was determined using a Kruskal–Wallis test.

**Fig. S2. Extended analyses of in vivo engraftment of "trained" APL blasts.**

(A) Representative picture of human peripheral blood-derived M2d macrophages post-polarization (left picture) and after 48h co-culture with human primary APL cells (right picture). (B) Bar graph showing the cell counts of primary APL cells when put in co-culture with macrophages at the day 0 (initial input) and after 2 days of co-culture. D0 = Day 0, D2 = Day 2. (C) Dispersion graph showing the leukocyte count over time of mice injected with human primary APL cells pre-cultured on bone marrow-derived human primary mesenchymal stromal cells (MSCs) for 48h. (D) Human $CD45^+$ and $CD33^+$ chimerism level detected in different organs at week 18 post-transplant of 22 mice injected with human primary APL cells pre-cultured on primary MSCs (Passage 4) for 48h. (E) Human $CD117^+CD33^+$ chimerism levels (%) measured in the spleen of secondary transplanted mice. Representative cytospins. (F) Bar graph showing the human CD45 chimerism level measured in the blood of mice transplanted with $1x10^3$ sorted human $CD33^+CD117^+$ APL blasts from M2d pre-culture primary transplant. (G) Dot plot graph showing the murine CD45.2 chimerism level of mice transplanted with murine primary APL blast cells (from hCG-PML-RARA mice – $CD45.2^+$) alone (control) or pre-cultured on either M0 or M2d murine macrophages inside sub-lethally irradiated (350 cGy) C57BL/6J.PepBoy recipients ($CD45.1^+$). (H) Overall survival analysis of C57BL/6J transplanted with murine primary APL blast cells alone or precultured on either M0 or M2 murine macrophages. (I) Murine APL blast cells defined by murine $CD117^+$ and $Gr1^{+/int}$ detected in the BM of mice transplanted with murine primary APL blast cells alone or pre-cultured on either M0 or M2d murine macrophages post-euthanasia. (J) Bar graph showing the frequency of early ($CD117^+Gr1^+CD34^+CD16/32^-$) and late promyelocytes ($CD117^+Gr1^+CD34^-CD16/32^+$) of mice transplanted with murine primary APL blast cells alone or pre-cultured on either M0 or M2d murine macrophages post-sacrifice. Bellow, representative bone marrow smears with the cells present in the BM of mice transplanted with APL blasts alone (APL control) and co-cultured with M2d-macrophages (APL M2d co-culture).

(B–D) Each dot represents an individual APL patient

(G-J) Each dot represents an individual mouse

(B) Wilcoxon signed rank test (2-sided) *$P<0.05$.

(H) Survival curves were plotted using the Kaplan-Meier method and compared by the log-rank test.

One-way (E and I-J) or two-way (C, F and G) analysis of variance (ANOVA).

**Fig. S3. Extended data on mechanisms involved in M2d-mediated reprogramming of primary AML cells.**

(A) Bar graph showing the level of primary AML phagocytosis at diagnosis (day 0) on non-polarized (M0) macrophages. Representative pictures of low and high phagocytosed AML cells. Macrophages are labeled in red (CD11b-AF594) and tumor cells in green (CellTrace™ CFSE Cell proliferation AF488). (B) Pearson correlation graphs between the level of phagocytosis measured at diagnosis and the CD47 and CD24 MFI level as well as the percentage of CD24 and Calreticulin (CALR) measured on the respective AML blast population ($SSC^{low}CD45^{dim}CD34^+$ or $CD117^+$ for $CD34^-$ AMLs). MFI= Mean fluorescence intensity, CALR=Calreticulin. (C) Bar graph showing the MFI of CD47/CD24 and CD24 percentage at diagnosis and after two-day co-culture on M2d macrophages. (D) Gene expression of STING related genes, measured in primary M1 and M2d-macrophage samples at diagnosis and after a 48h co-culture with primary AML blasts. Data is plotted as relative expression to *RPL30* and *ACTB* endogenous controls (E) Dot plot graph showing the fold change to diagnosis of APL/AML cells migrated post M2d macrophage exposure for 48h. Migration assay was performed using transwell system, with healthy human MSCs plated in the lower chamber (Passage 4). (F) MFI levels of CD49d and CD49f measured in primary AML and APL blasts ($CD45^+CD34^+$ for AML samples and $CD45^+CD117^+$ for APL samples) cells at diagnosis and after a 2-day co-culture on M2d macrophages. (G) Percentage of CD49d measured in human APL and AML blasts ($CD45^+CD117^+CD33^+$/

$CD45^+CD33^+$) cells post-sacrifice of primary and secondary APL xenotransplant (left and middle panel, respectively) and primary AML xenograft (right panel).

(A) Each dot represents a different counted field

(B-E) Each dot represents an independent AML patient.

(C-E) Wilcoxon signed rank test (2-sided) *P<0.05.

One-way (F) analysis of variance (ANOVA).

**Fig. S4. Extended data on mitochondrial exchange form macrophages to leukemic blasts.**

(A) Bar graph of basal and maximum oxygen consumption rate (OCR) of HL60 cells after two-day co-culture on MS5 cells and M2d macrophages. (B) Bar graph of OCR and extracellular acidification rate (ECAR) of primary AMLs after two-day co-culture on MS5 cells and M2d macrophages. (C) Mitochondrial transfer measured in HL60 and MOLM13 cells after co-culture on mitochondrial labeled MS5 cells and M2d macrophages. MS5 and M2d macrophages were labeled with MitoTracker Green™ (2 μM) and DeepRed™ (1 μM) for 15 min and incubated for 48 hours to remove potential dye leakage. Conditioned medium (CM) of stained macrophages were used as negative control of unspecific staining. Mitochondrial transfer was determined by measuring the MitoTracker in the AML cell lines (Gating inside the $CD45^+$ cells for the MS5 co-cultures or HLA-DR- cells for the M2d co-cultures). Representative histogram showing the mitochondrial mass (MitoTracker Green$^{TM}$) and potential (MitoTracker DeepRed$^{TM}$) (Right panels). (D) Evaluation by real-time quantitative polymerase chain reaction of murine mitochondrial DNA content on AML cell lines (MOLM13 and HL60) and primary AML samples after 48h of co-cultures with MS5 and M2d macrophages. Values are normalized to MS5 control cells. (E) Colony formation of primary AML blasts (methylcellulose) scored after 14 days.

(A-B) Each dot represents an independent biological replicate (HL60) or AML patient. Values represent the average of 4 technical replicates for each experiment.

(E) Each dot represents an independent experiment

25

(C-D) Represent at least 3 biological replicates for all experiments (performed with 4 technical replicates, each).

(A-B and E) Wilcoxon signed rank test (2-sided) **P<0.01.

**Fig. S5. Extended data on the effect of macrophages after a two-day co-culture and when macrophages are treated with etomoxir.**

(A) Basal, maximum OCR and ECAR of M2d macrophages treated with vehicle (DMSO) or Etomoxir (50 µM) for 24 hours. (B) Bar plot comparing the levels (%, upper panel; MFI, lower panel) of M2- (CD163 and CD206) and M1- (CD80 and CD86) markers measured by FACS in healthy activated M2a- and M2d-macrophages treated with vehicle and Etomoxir (50 µM) for 24 h. (C) Maximum OCR of HL60 cells cultured on either MS5 or M2d cells pretreated with vehicle or etomoxir for 24h. Two-way (B and C) analysis of variance (ANOVA). (D) Cumulative cell count in liquid culture of primary AML cells exposed to MS5 or M2d macrophages treated with vehicle or etomoxir (50 µM). (A) Wilcoxon signed rank test (2-sided). *P<0.05.

**Fig. S6. Extended data on the M2-UMCG signature.**

(A) Levels of M1 and M2 macrophages measured by QuanTIseq according to the M2-UMCG signature groups in the TCGA cohort. (B) Kaplan-Meier (KM) analysis of 26 OS using the M2-UMCG signature in the BeatAML cohort. (C) Levels of M1 and M2 macrophages measured by QuanTIseq according to the M2-UMCG signature groups in the BeatAML cohort. (D) Levels of M1 and M2 macrophages measured by QuanTIseq according to the M2-UMCG signature groups in the MDS patient cohort. (E) Distribution of the M2-UMCG signature across all IPSS-R classification groups.

(A and C-D) Mann-Whitney test. **P <0.01, ***P <0.001, NS, not significant.

(B) Patients were dichotomized into low and high M2-UMCG signature. Survival curves were estimated using the KM method, and the log-rank test was used for comparison.

(E) Chi-square test

### Figure S1

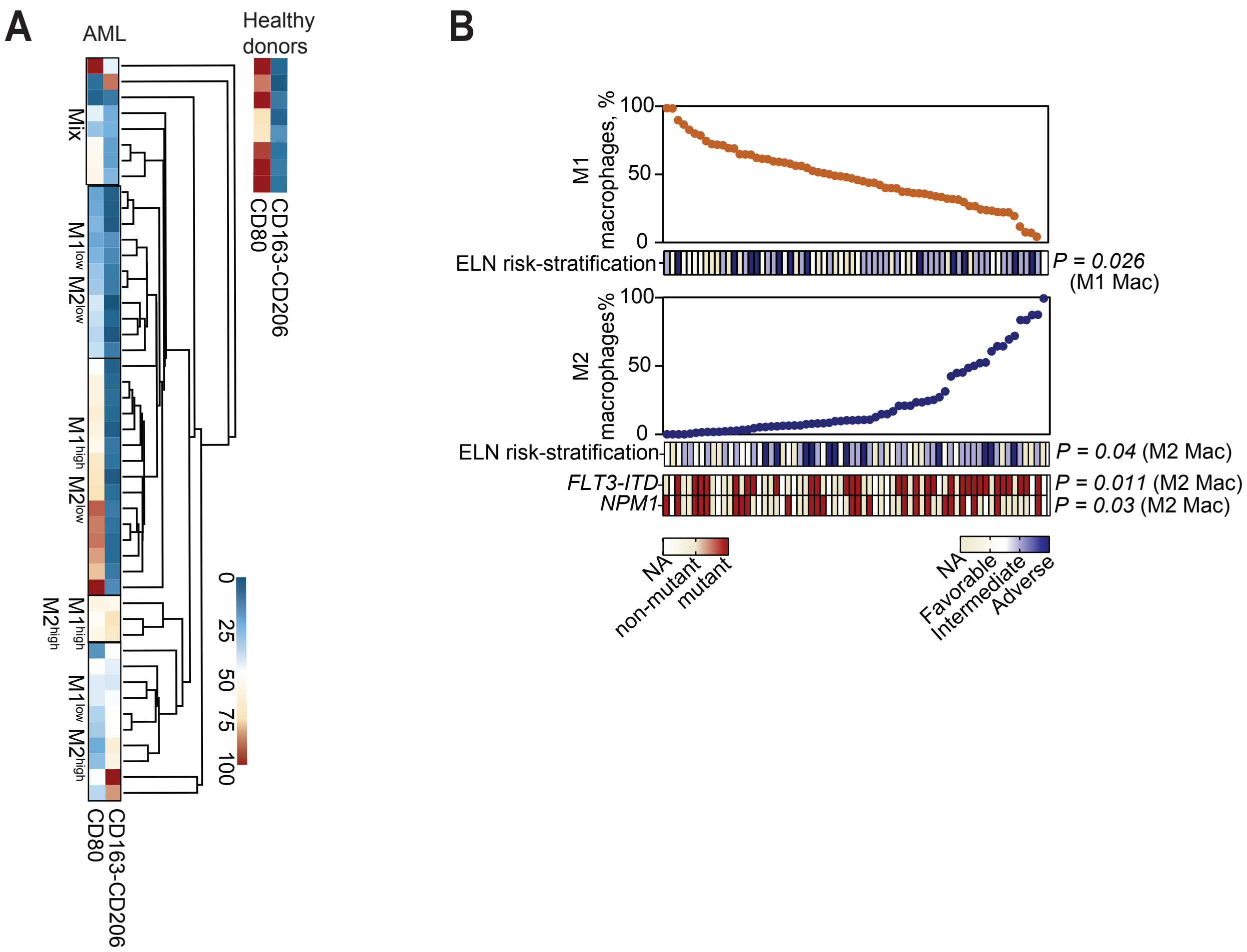
A
AML
Healthy donors
Mix
M1low M2low
M1high M2low
M1high M2high
M1low M2high
CD80
CD163-CD206
0
25
50
75
100
B
M1 macrophages, %
100
50
0
ELN risk-stratification
P = 0.026 (M1 Mac)
M2 macrophages%
P = 0.04 (M2 Mac)
FLT3-ITD
P = 0.011 (M2 Mac)
NPM1
P = 0.03 (M2 Mac)
NA
non-mutant
mutant
Favorable
Intermediate
Adverse

Figure S2

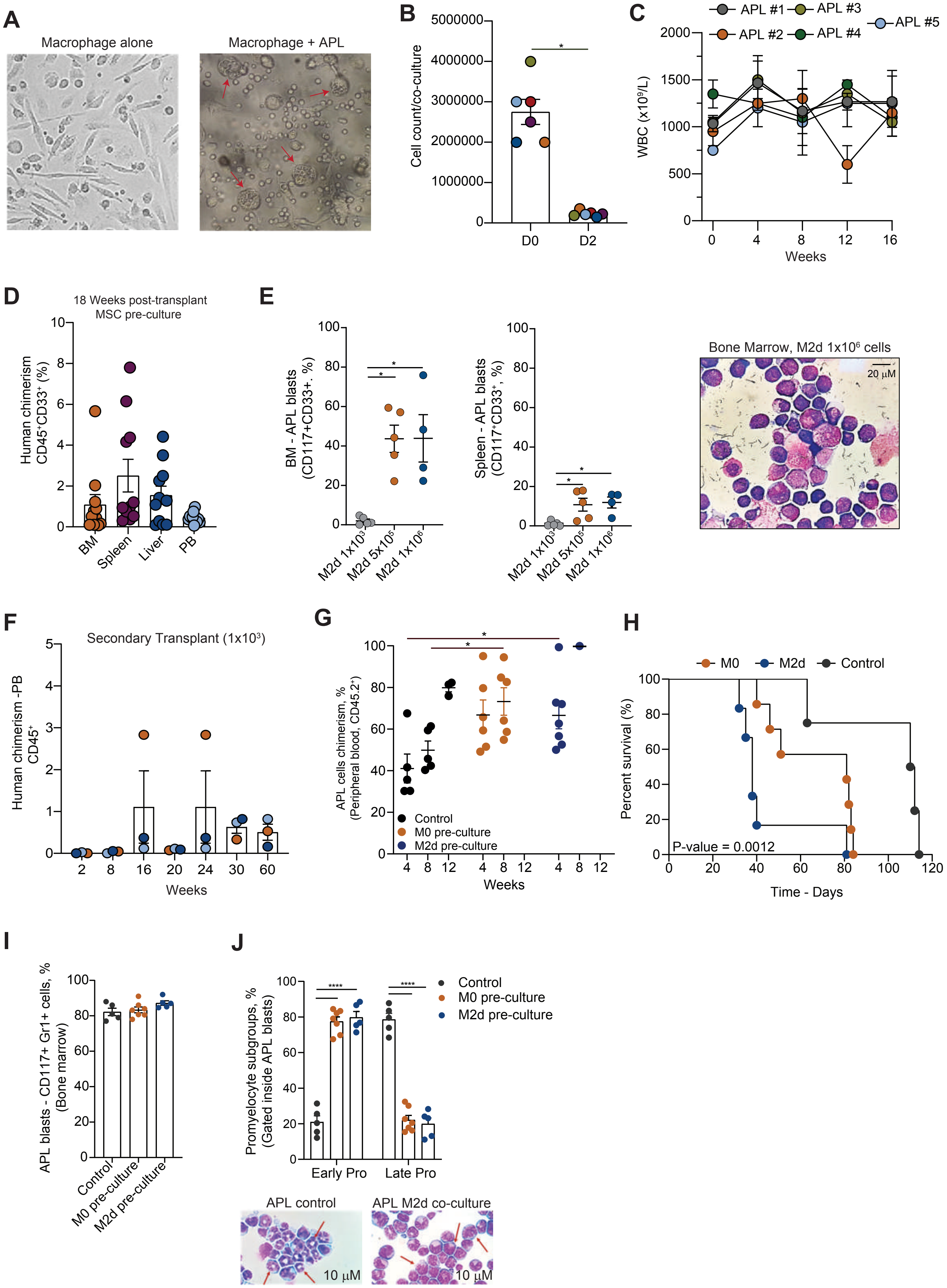
A
Macrophage alone
Macrophage + APL
B
Cell count/co-culture
D0
D2
C
MSC Pre-culture
APL #1
APL #2
APL #3
APL #4
APL #5
WBC (x10⁹/L)
Weeks
D
18 Weeks post-transplant
MSC pre-culture
Human chimerism CD45⁺CD33⁺ (%)
BM
Spleen
Liver
PB
E
BM - APL blasts (CD117+CD33+. %)
Spleen - APL blasts (CD117+CD33⁺, %)
M2d 1x10³
M2d 5x10⁵
M2d 1x10⁶
Bone Marrow, M2d 1x10⁶ cells
20 µM
F
Secondary Transplant (1x10³)
Human chimerism -PB CD45⁺
Weeks
G
APL cells chimerism, % (Peripheral blood, CD45.2⁺)
Control
M0 pre-culture
M2d pre-culture
Weeks
H
M0
M2d
Control
Percent survival (%)
P-value = 0.0012
Time - Days
I
APL blasts - CD117+ Gr1+ cells, % (Bone marrow)
Control
M0 pre-culture
M2d pre-culture
J
Promyelocyte subgroups, % (Gated inside APL blasts)
Control
M0 pre-culture
M2d pre-culture
Early Pro
Late Pro
APL control
APL M2d co-culture
10 µM
10 µM

Figure S3

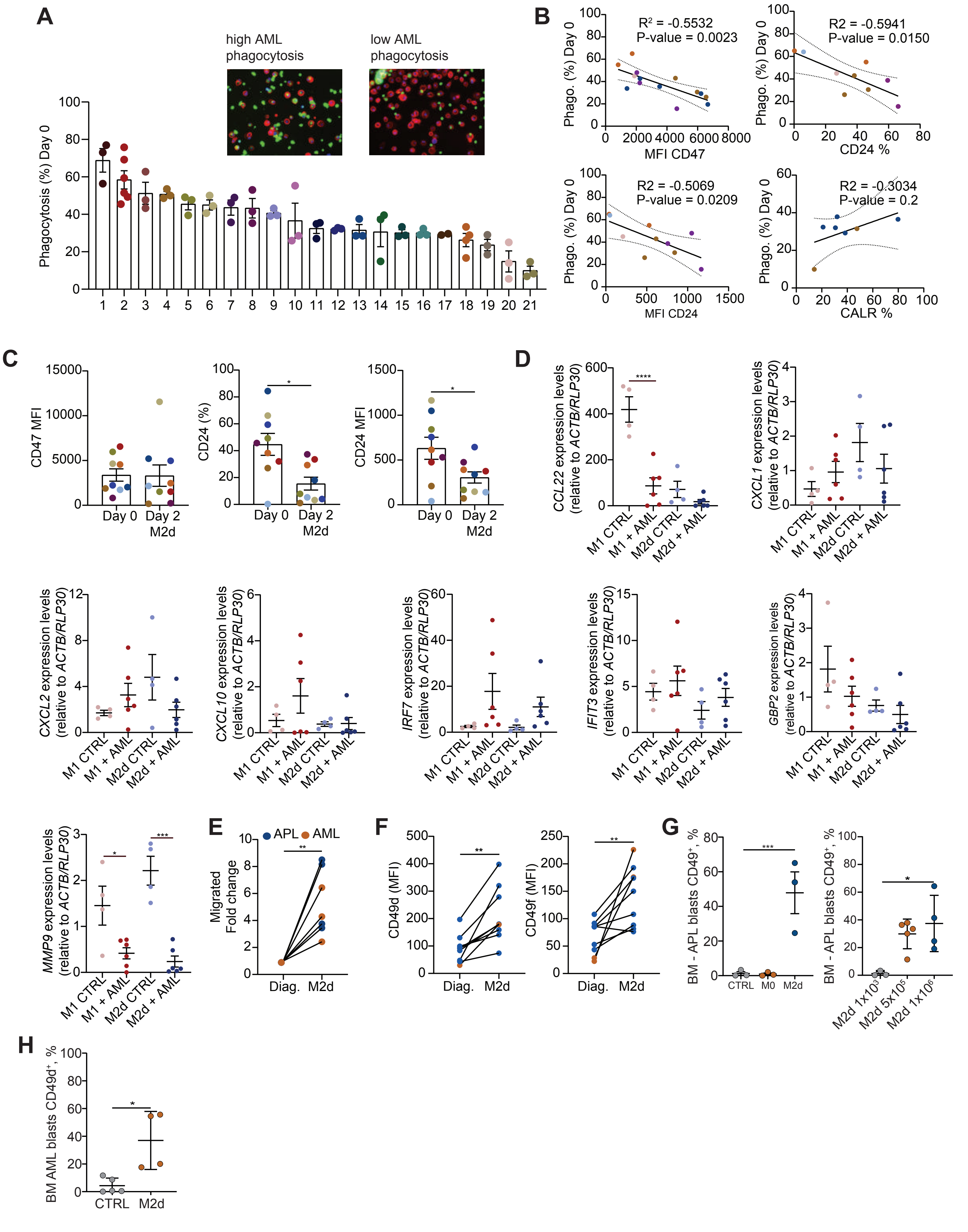

A
high AML phagocytosis
low AML phagocytosis
Phagocytosis (%) Day 0
B
R² = -0.5532
P-value = 0.0023
Phago. (%) Day 0
MFI CD47
R2 = -0.5941
P-value = 0.0150
CD24 %
R2 = -0.5069
P-value = 0.0209
MFI CD24
R2 = -0.3034
P-value = 0.2
CALR %
C
CD47 MFI
CD24 (%)
CD24 MFI
Day 0
Day 2 M2d
D
CCL22 expression levels (relative to ACTB/RLP30)
CXCL1 expression levels (relative to ACTB/RLP30)
CXCL2 expression levels (relative to ACTB/RLP30)
CXCL10 expression levels (relative to ACTB/RLP30)
IRF7 expression levels (relative to ACTB/RLP30)
IFIT3 expression levels (relative to ACTB/RLP30)
GBP2 expression levels (relative to ACTB/RLP30)
MMP9 expression levels (relative to ACTB/RLP30)
M1 CTRL
M1 + AML
M2d CTRL
M2d + AML
E
APL
AML
Migrated Fold change
Diag.
M2d
F
CD49d (MFI)
CD49f (MFI)
G
BM - APL blasts CD49⁺, %
CTRL
M0
M2d
M2d 1x10³
M2d 5x10⁵
M2d 1x10⁶
H
BM AML blasts CD49d⁺, %
CTRL
M2d

Figure S4

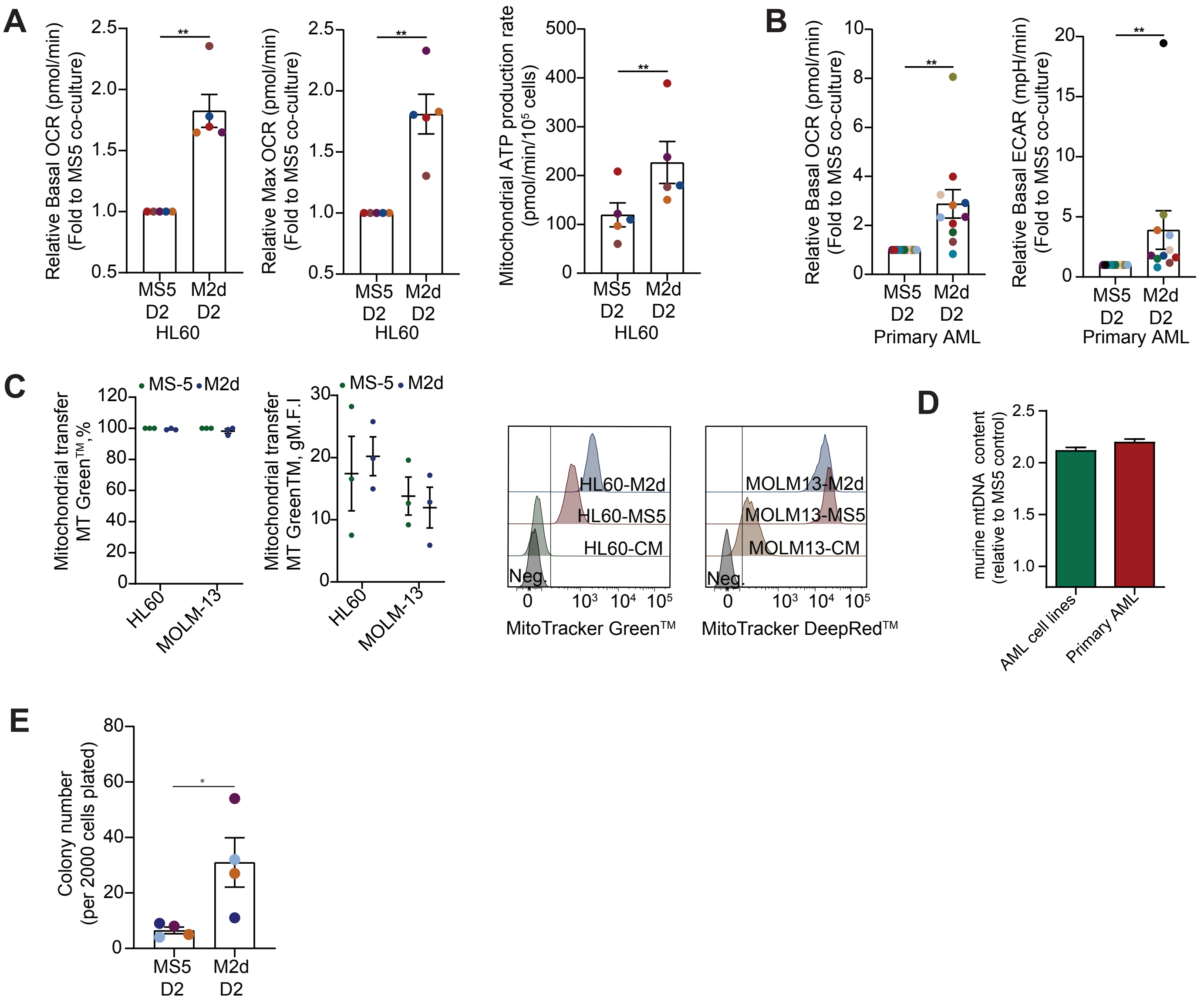
A
Relative Basal OCR (pmol/min)
(Fold to MS5 co-culture)
**
MS5
D2
M2d
D2
HL60
Relative Max OCR (pmol/min)
(Fold to MS5 co-culture)
Mitochondrial ATP production rate
(pmol/min/10⁵ cells)
B
Relative Basal OCR (pmol/min)
(Fold to MS5 co-culture)
Primary AML
Relative Basal ECAR (mpH/min)
(Fold to MS5 co-culture)
C
MS-5
M2d
Mitochondrial transfer
MT Green™,%
HL60
MOLM-13
Mitochondrial transfer
MT GreenTM, gM.F.I
HL60-M2d
HL60-MS5
HL60-CM
Neg.
MitoTracker Green™
MOLM13-M2d
MOLM13-MS5
MOLM13-CM
MitoTracker DeepRed™
D
murine mtDNA content
(relative to MS5 control)
AML cell lines
Primary AML
E
Colony number
(per 2000 cells plated)
*

Figure S5

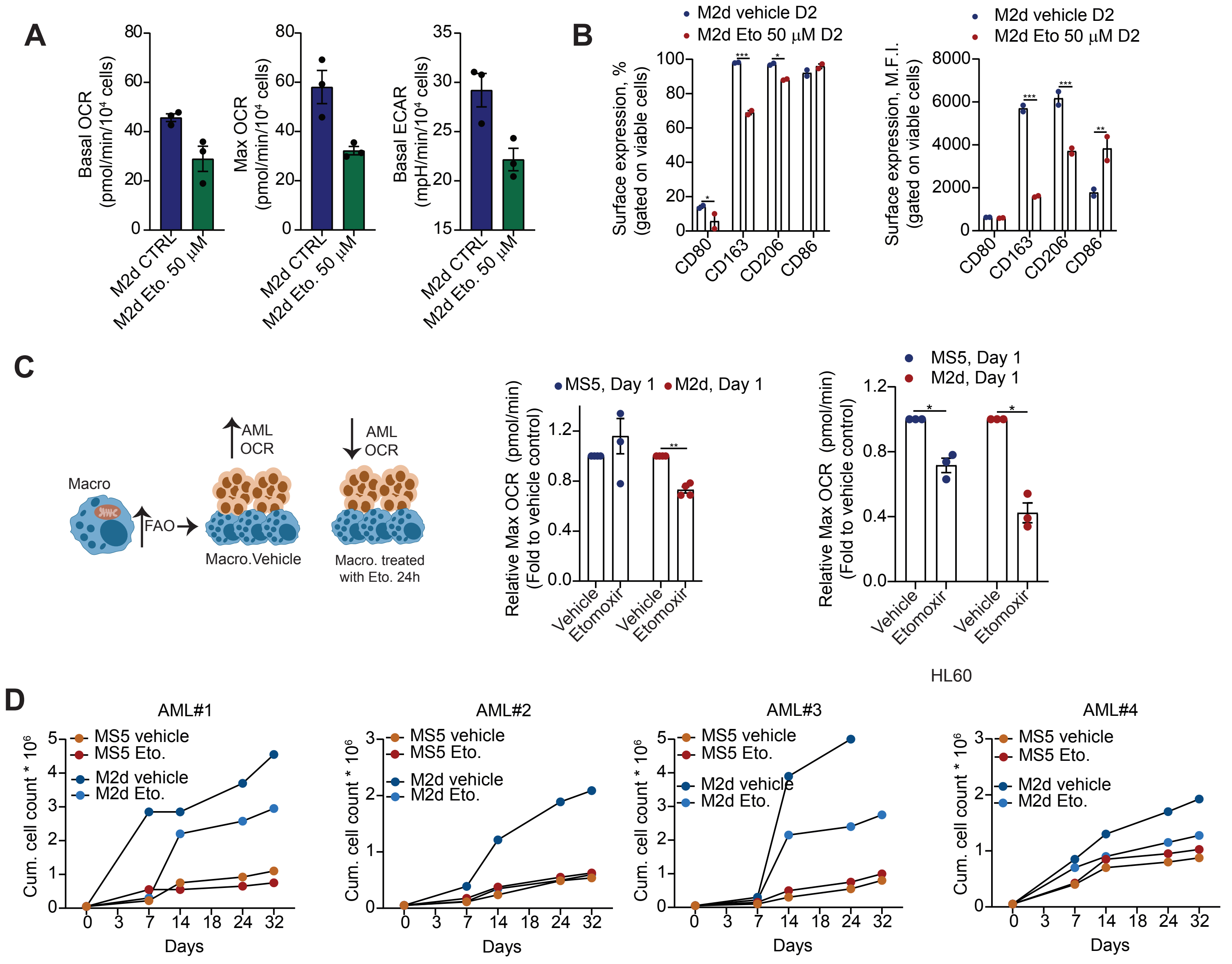
A
Basal OCR (pmol/min/10⁴ cells)
Max OCR (pmol/min/10⁴ cells)
Basal ECAR (mpH/min/10⁴ cells)
M2d CTRL
M2d Eto. 50 µM
B
M2d vehicle D2
M2d Eto 50 µM D2
Surface expression, % (gated on viable cells)
Surface expression, M.F.I. (gated on viable cells)
CD80
CD163
CD206
CD86
C
Macro
FAO
AML OCR
Macro.Vehicle
Macro. treated with Eto. 24h
MS5, Day 1
M2d, Day 1
Relative Max OCR (pmol/min) (Fold to vehicle control)
Vehicle
Etomoxir
HL60
D
AML#1
AML#2
AML#3
AML#4
MS5 vehicle
MS5 Eto.
M2d vehicle
M2d Eto.
Cum. cell count * 10⁶
Days

Figure S6

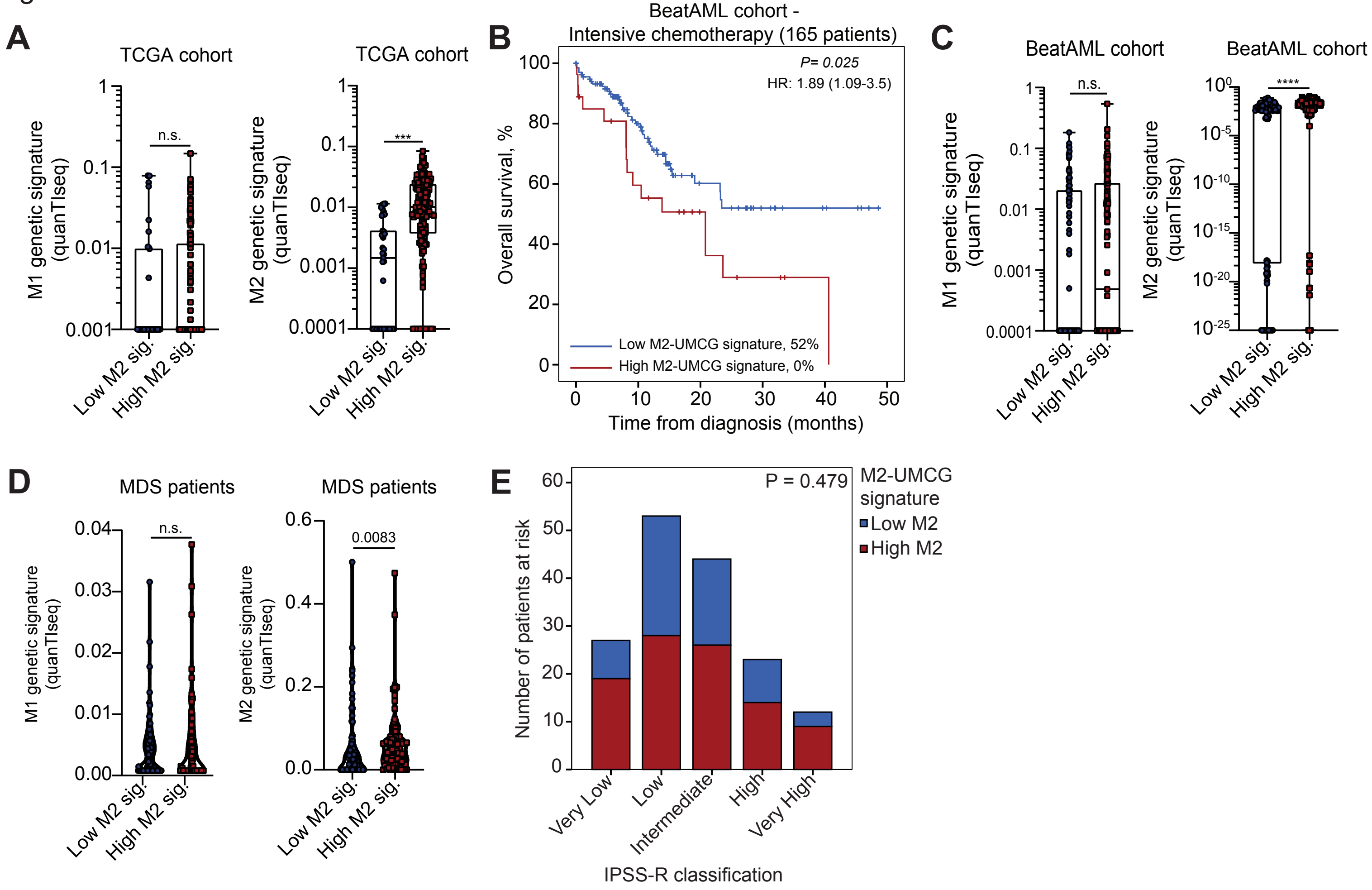

A
TCGA cohort
M1 genetic signature (quanTIseq)
n.s.
Low M2 sig.
High M2 sig.
TCGA cohort
M2 genetic signature (quanTIseq)
***
B
BeatAML cohort - Intensive chemotherapy (165 patients)
P= 0.025
HR: 1.89 (1.09-3.5)
Overall survival, %
Time from diagnosis (months)
Low M2-UMCG signature, 52%
High M2-UMCG signature, 0%
C
BeatAML cohort
M1 genetic signature (quanTIseq)
n.s.
BeatAML cohort
M2 genetic signature (quanTIseq)
****
D
MDS patients
M1 genetic signature (quanTIseq)
n.s.
MDS patients
M2 genetic signature (quanTIseq)
0.0083
E
P = 0.479
Number of patients at risk
M2-UMCG signature
Low M2
High M2
Very Low
Low
Intermediate
High
Very High
IPSS-R classification
